# Telomemore enables single-cell analysis of cell cycle and chromatin condensation

**DOI:** 10.1101/2023.03.19.533267

**Authors:** Iryna Yakovenko, Ionut Sebastian Mihai, Martin Selinger, William Rosenbaum, Andy Dernstedt, Remigius Gröning, Johan Trygg, Laura Carroll, Mattias Forsell, Johan Henriksson

## Abstract

**Graphical abstract:** 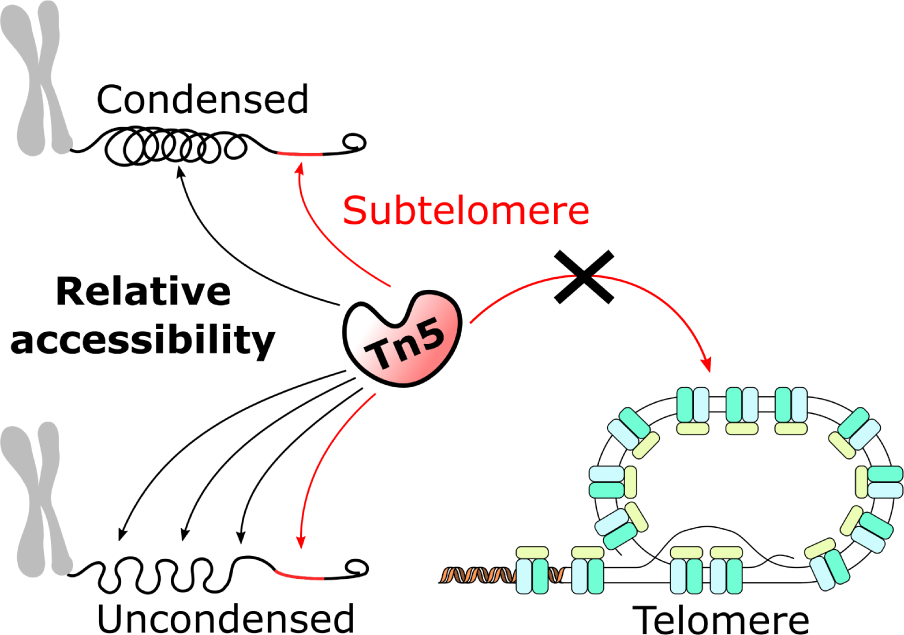

**ABSTRACT:** Single-cell RNA-seq methods can be used to delineate cell types and states at unprecedented resolution but do little to explain why certain genes are expressed. Single-cell ATAC-seq and multiome (ATAC+RNA) have emerged to give a complementary view of the cell state. It is however unclear what additional information can be extracted from ATAC-seq data besides transcription factor binding sites. Here we show that ATAC-seq telomere-like reads, mostly originating from the subtelomere, cannot be used to infer telomere length, but can be used as a biomarker for chromatin condensation. Using long-read sequencing, we further show that modern hyperactive Tn5 does not duplicate 9bp of its target sequence, contrary to common belief. We provide a new tool, Telomemore, which can quantify non-aligning subtelomeric reads. By analyzing several public datasets, and generating new multiome fibroblast and B cell atlases, we show how this new readout can aid single-cell data interpretation. We show how drivers of condensation processes can be inferred, and how it complements common RNA-seq-based cell cycle inference, which fails for monocytes. Telomemore-based analysis of the condensation state is thus a valuable complement to the single-cell analysis toolbox.

## INTRODUCTION

Maintenance of genome integrity is an essential biological process, as errors can lead to loss of genes or cancer. The end of chromosomes (telomeres) need special treatment, as the typical DNA replication process results in the loss of 50-100bp per cell division (1). To overcome this, various mechanisms exist to extend the telomere, which in e.g. humans and mice, mainly consists of 6bp tandem repeats of TTAGGG (Figure 1a), protected by the Shelterin complex (2). Because short telomeres can indicate or be the cause of over 13 pathogenic conditions (from liver fibrosis to cancer (3)), there is considerable interest in measuring the length of this region. Several methods, such as qPCR, Terminal restriction fragments Southern blot (TRF), fluorescence *in situ* hybridization (FISH), and long-read sequencing have been developed (4, 5). There is however a need to study telomere length also in individual cells, and correlate it with other features of interest. In cancer cells, for example, telomeres can be both shorter and longer than in normal cells (6); such single-cell differences may not be detected if measured using “bulk” methods, which provide an average over many cells. Furthermore, telomere length varies across cell types, and without an understanding of what cell is being studied, the length may have little meaning. As a result, telomere length from bulk cancer measurement is not considered a good biomarker (4).

Single cell RNA-seq is the current state-of-the-art tool to understand heterogeneity of cells in a tissue. Limited efforts have been made to also include telomere length measurement, but the largest study to date only measured telomere lengths in 242 cells, likely because it was performed in low-throughput and costly multi-well plates (7). As contemporary RNA-seq-only single-cell atlases can easily measure 100k -1M cells, this begs for a new single-cell approach to simultaneously measure cell state and telomere length.

ATAC-seq (Assay for Transposase-Accessible Chromatin using sequencing) is a widely used method for characterizing the epigenetic state by finding open chromatin regions (8). ATAC-seq is based on the enzyme Tn5 fragmenting and tagging (“tagmenting”) open chromatin, which is then sequenced (Figure 1b). The accessibility of regions can then be compared, and in particular, transcription factor (TF) activity can be quantified based on motif presence in accessible regions (9). Thus, potential upstream regulators of different cell types and fates can then be pinpointed. Gene expression levels can also be approximated from the accessibility of their transcriptomic start site (TSS). With the goal of generating novel readouts from ATAC-seq data, we discovered telomere-like reads. We hypothesized that potential tagmentation of telomeres could also be used for telomere length measurement, as longer telomeres could possibly result in more ATAC-seq reads. Thus, we analyzed both public and novel data to validate this hypothesis.

Here, we show that single-cell ATAC-seq is unsuitable for inferring telomere length, largely because the telomere proper appears protected from Tn5 transposition. Instead, we show that, in most cases, presence of telomere-like reads rather indicate general condensation of the chromatin, as the subtelomere becomes relatively more accessible during condensation (Figure 3i). While the cell cycle stage can be inferred from RNA-seq, we claim that ATAC-seq sometimes is a more direct assessment of the entry to, and exit from, mitosis. Further analysis of the centromere may also help separate between early and late G_1_ phase. We provide a tool (Telomemore, https://github.com/henriksson-lab/telomemore.java) to assess chromatin condensation. This is validated using a new long-read atlas of Tn5 insertions, as well as by a convolutional neural network trained to approximately infer the origin of short reads. We provide examples, which showcase how metrics reported by Telomemore can aid in the interpretation of single-cell datasets, including (1) an analysis of transcription factors controlling the cell cycle, as well as (2) how CDKN1C seems to lock atypical monocytes in G_1_ phase. Finally, we provide a new single-cell atlas of human tonsil B cells and show that somatic hypermutation can be contextualized in terms of chromatin condensation. By providing precomputed chromatin condensation scores for several public single-cell datasets, we envision that further insights into the regulation of overall chromatin state will be unlocked.

## MATERIALS AND METHODS

### Acquisition of public single-cell data

Suitable datasets for reanalysis were mainly found from the 10x Genomics publication database (https://www.10xgenomics.com/datasets). To determine if sequencing reads were available, datasets were manually inspected using the National Center for Biotechnology Information (NCBI) Sequence Read Archive (SRA) online preview function (https://www.ncbi.nlm.nih.gov/sra). Whenever datasets had missing cell barcodes (“technical reads”), we instead attempted to obtain the complete original data using the SRA cloud retrieval function. We note that in SRA, cell barcode reads are inconsistently denoted as “biological” or “technical”. To ensure that we obtained the cell barcodes, we retrieved the data using the command “fasterq-dump SRRxxx -e 10 -v --include-technical --split-files”.

Furthermore we processed and broadly analyzed the following datasets: Delacher2021 (10) (GSE156112), Jain2021_pbmc (11) (E-MTAB-11225, E-MTAB-11226), Jain2021_thymus (11) (E-MTAB-9828, E-MTAB-9840), Lyu2021 (12) (GSE183684), Morabito2021 (13) (GSE174367), Sarropoulos2021 (14) (E-MTAB-9765), Satpathy2019 (15) (GSE129785), Taavitsainen2021 (16) (GSE168667), Wimmers2021 (17) (GSE165904), Ziffra2021 (18) (GSE163018), Kinoshita2021 (19) (GSE131549), Qiangli2021 (20) (GSE178551) and Zhang2021 (21) (GSE184462).

The Satpathy2019 GEO dataset GSE129785 was not compatible with ArchR. Also, as cell barcodes were missing in the SRA upload, SRA cloud delivery was used. BAM files were converted to FASTQ using bamtofastq (https://github.com/10XGenomics/bamtofastq/), and then realigned using CellRanger. Similarly, we solved issues with the download of Ziffra2021 and Delacher2021 using the cloud delivery service.

For multiome mouse embryo data (Argelaguet2022) (22), rather than realign the raw files (GSE205117), we used the processed count and ATAC fragment files provided via FTP (https://github.com/rargelaguet/mouse_organogenesis_10x_multiome_publication).

### Generation of a tonsillar B cell atlas

The research was carried out according to The Code of Ethics of the World Medical Association (Declaration of Helsinki). Ethical permits were obtained from the Swedish Ethical review authority (No: 2016/53-31), and all samples were collected after receiving informed consent from patients or patients’ guardians. Briefly, tonsillar cell suspensions were prepared by tissue homogenization in RPMI-1640 medium and passed through a 70 µm cell strainer. Red blood cells were lysed using BD PharmLyse lysis buffer according to the manufacturer’s instructions. All cell suspensions were frozen in fetal bovine serum (FBS) (Gibco) with 10% DMSO and stored in liquid N_2_.

B cells from tonsils of 5 individuals were enriched using negative selection magnetic beads (EasySep Human B Cell Isolation kit, Stemcell technologies, #17954). They were then pooled to avoid batch effects, with an average viability per donor of 92% (S.D. 1.5%).

Enriched B cells were washed in ice-cold ATAC-seq resuspension buffer (RSB, 10 mM Tris pH 7.4, 10 mM NaCl, 3 mM MgCl_2_), spun down, and resuspended in 100 mL ATAC-seq lysis buffer (RSB plus 0.1% NP-40 and 0.1% Tween-20, Thermo Fisher). Lysis was allowed to proceed on ice for 5 min, then 900 mL RSB was added before spinning down again and resuspending in 50 mL 1X Nuclei Resuspension Buffer (10x Genomics). To assess nuclei purity and integrity after lysis, nuclei were stained with Trypan Blue (15250061, Thermo Fisher Scientific), and DAPI (D1306, Thermo Fisher Scientific), according to the manufacturer’s recommendation. If necessary, cell concentrations were adjusted to equal ratios (per donor) prior to starting single-cell GEM emulsion droplet generation with the ATAC-seq NextGEM kit (10x Genomics). Briefly, nuclei were incubated in a transposition mix. Transposed nuclei were then loaded into a Chromium Next GEM Chip J. A total of 9000 nuclei were loaded per lane, with a target recovery of 5500 (doublet rate 4% - 4.8%). After generation of GEM emulsions, we performed reverse transcription, cDNA amplification and library indexing according to manufacturer specifications (*Chromium Next GEM Single Cell Multiome ATAC + Gene Expression*, CG000338 Rev A, 10x Genomics, September 2020).

### Generation of a colonic fibroblast multiome atlas

Human colonic fibroblasts (CRL-1459, ATCC) were maintained in Eagle’s Minimum Essential Medium (EMEM; 30-2003, ATCC) supplemented with 10% of FBS (10270-106, Gibco, ThermoFisher Scientific) and 10 μg/ml of ciprofloxacin (17850, Sigma-Aldrich Merck), as described before (23). A single-cell multiome library was generated analogously to the B cell libraries, with lysis continuing for 5 minutes, and 3000 nuclei loaded into the Chromium instrument. Passage 10 and 15 were collected, but only passage 10 was included here due to (1) large differences in telomere-like read abundances, and (2) the latter passage seemed to enter terminal differentiation, and the heterogeneity was too complex for the purposes of our validation. Raw data have been submitted to ArrayExpress.

### Telomere *k*-mer motif data analysis in short-read data

The Telomemore pipeline was first implemented in Python, and later in Java for performance and for the ease of supporting multiple input file formats in an object oriented manner. To detect telomeric reads, CCCTAA and TTAGGG-motif scanning was performed, either using (1) custom python scripts with FASTQ files as input, or (2) using the Telomemore software, which also supports BAM files as input and cell barcode assignments. At least 3 consecutive telomere motifs were required for a read to be counted, which was qualitatively motivated based on a histogram analysis, but has also been motivated previously based on comparison to the gold standard Terminal restriction fragments Southern blot (TRF) (24).

To be able to compare the number of telomeric reads between libraries, we defined nTA (normalized telomere availability) as the number of telomere-like fragments divided by the total number of reads:

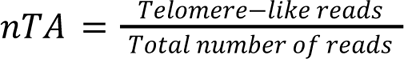

This measure takes sequencing depth into account (for a whole bulk ATAC library, or per single cell). For visualization, we used rank(nTA) to maximize contrast.

### Analysis of bulk aging PBMC ATAC-seq

We aligned the PBMC *vs* aging dataset (25) to the cellranger-arc-GRCh38-2020-A-2.0.0 reference genome using STAR v2.7.1.a. To find sources explaining the variation, we calculated the nucleosome signal from the histogram of mapped read lengths, computed by ATACFragQC v0.4.5. Furthermore, we qualitatively investigated all metadata available for each donor (e.g., by considering read length, as one batch in particular had a different read length; by considering telomere content, as telomere content was dramatically different for donors with high donor IDs; by assessing variant telomere repeats [T**GGG]). Overall we identified 4 batches A-D; Batch D is samples with subject ID > 260, C remainder with average read length > 130, B remainder with read length > 80, and batch A contains the rest of the samples. We then fitted telomere abundance *vs* age for each batch. We also attempted several variants of multilinear models, including covariates such as sex, nucleosome signal, batch, and number of centromeric reads (inferred by CNN model, described later). We further attempted to include the number of variant telomere repeats (TAAGGG, TTTGGG, etc), inspired by previous work (26). None of the aforementioned approaches resulted in a statistically significant or consistent correlation with age across batches.

The bulk PBMC dataset was also used to study the binding preference motif of Tn5. Since short telomeric reads cannot reliably be aligned to the reference genome, a reference-free approach was used to reorient the reads (Suppemental Figure 1d). In short, if the G% content was below 50% on the 5’ side, the forward (R1) and reverse (R2) reads were selectively reverse complemented and swapped. A custom R script was then used to estimate base frequencies at each position.

### Inference of the approximate origin of reads using deep learning

A convolutional neural network was implemented in PyTorch with the aim of predicting the average position of a 50bp sequence from the telomere. The distance is defined as 0 for any sequence within the telomere proper. The centromere region was taken from the T2T reference genome using the associated repeat masker GTF-file, while the chromatin proper is defined as the remainder, up to 10 Mbp from the end of each chromosome. The sequence was then extracted using getfasta from Bedtools 2.30.0 (27).

To ensure that the natural human variability of the subtelomere is handled by the algorithm, nanopore data of the telomeres of 18 individuals was obtained (28, 29) (IDs in dataset: 3048691-3048709). The sequence of the biotin adapter (CCCTCCGATA) ligated to the telomeres was trimmed from the beginning of each read. BAM files were read by invoking “samtools view” in a pipe using the Python subprocess library. Reads shorter than 8kb were omitted. To define the end of the telomere proper, the read sequence was repeatedly scanned for TAACCC within 0 to 18bp from the current position. When no more matches are found, the current position is set at the start of the subtelomere. The results were deemed similar to manual curation.

The philosophy of the network was to have few parameters to try and avoid overfitting, but still acceptable performance for the purpose. Each base of ATCGN was one-hot encoded for the input. Several designs were qualitatively evaluated. The final design was: a convolutional layer #1 (kernel size 6, 20 filters), ReLU, convolutional layer #2 (kernel size 6, 20 filters), ReLU, and a dense layer. The kernel size was selected to aid the network in detecting the telomere hexamer sequence. 500, 000 random centromere/chromatin sequences were used for training, along with 500, 000 random sequences from the telomere nanopore data. The reverse complements of the sequences, computed using Biopython (30), were also included. The data was split 80/20 into training and test datasets. The training error was defined as the mean squared error of the predicted log(distance) from the telomere. The model was trained for 200 epochs although 100 epochs seemed sufficient. No overfitting was noticed based on test data predictions.

The precise log(distance) prediction was not used for comparison with nTA, as further hyperparameter optimization and deep validation is likely required. Instead, predictions were binned to be within the distances of each respective training dataset, and reads with distance=0 were considered subtelomeric (as such reads are very sparse and long-read data suggest them to seldom be from the telomere proper).

### Analysis of TCGA cancer data

All 410 ATAC-seq libraries were downloaded from TCGA, effectively from a single study (31). Using the TCGA portal, the corresponding whole-genome sequencing (WGS) samples were located. Telomere lengths were then obtained from a previous analysis (32) using TelSeq (24). Because ATAC nTA is affected by cell type, each cancer type was analyzed separately. Qualitatively, no correlation was found.

### Generation of long-read WGS libraries after Tn5 transposition

CCRF-CEM (CCL-119, ATCC) and Jurkat cells, both of T cell origin, were cultured in RPMI 1640 medium (cat no. 21870076, Thermo Fisher Scientific) with 10% Bovine Serum (cat no. A3294-10G, Sigma Aldrich) 1% 100× penicillin-streptomycin-glutamine. For transpososome assembly the preanealed cargo 250 nM duplex PAGE purified oligos were ordered from IDT (Coralville): sense (5’-CTGTCTCTTATACACATCTACGCGGTGGACAAAAAATTTCATTTGGAACTAGATTTGACCTCAGCTTCAATGCCAGAGATGTGTATAAGAGACAG) and antisense (5’-CTGTCTCTTATACACATCTCTGGCATTGAAGCTGAGGTCAAATCTAGTTCCAAATGAAATTTTTTGTCCACCGCGTAGATGTGTATAAGAGACAG). Cargo oligos of 95bp were created with Mosaic ends (ME) (33) and with a restriction site for BbvCI. Such length was previously found in one of the potential optimal windows for binding Tn5 transposase (34). The body of the cargo was modified from the parts of the Tn5 transposon (GenBank KF813062.1) that were not creating the hairpin according to the folding tool Mfold (35). The cargo concentration was adjusted to 67 µM in the buffer (2mM DTT, 771mM NaCl, 44.7mM Tris-HCl pH 7.5, 10% glycerol) and stored at -20°C. Tn5 _E54K, L372P_ manufactured and purified at EMBL Heidelberg (36) was removed from -80°C and thawed. Glycerol concentration was adjusted to 50% and stored for a few weeks at -20°C. Tn5 was assembled with cargo oligonucleotides at equal molar. The mixture was incubated at room temperature, shaking at 400 revolutions per minute (rpm) for 1 h. ATAC-seq (37) was modified to enable long-read analysis. Briefly, CCRF-CEM and Jurkat cells were grown overnight, and 100, 000 cells were were harvested at 500 × *g* 4 °C for 5 min. Cell pellets were resuspended in 50 μL of lysis buffer (10 mM Tris-HCl pH 7.4, 10 mM NaCl, 3 mM MgCl_2_, 0.1% NP-40, 0.1% Tween, 0.01% Digitonin) and kept for 4 min on ice, after which 500 μL of wash buffer (10 mM Tris-HCl pH 7.4, 10 mM NaCl, 3 mM MgCl_2_, 0.1% Tween) was added and gently resuspended by pipetting and centrifuged at 500 × *g* 4 °C for 15 min. The nuclei pellets were resuspended in the transposition reaction mix (TD buffer with 0.1 % Tween, 0.01% Digitonin, containing Tn5 transposase). The final concentrations of Tn5 in the reaction were 50 ng/µl. Reactions were incubated for 2 h at 37 °C shaking at 700 rpm. The transposed gDNA was purified using the Monarch Genomic DNA Purification Kit (catalog no. T3010S, New England Boilabs) and length and quality were accessed by gel. The estimated amount of inserts was assessed by qPCR with primers against the insert (F 5’-ACGCGGTGGACAAAAA, R 5’-CTCTTATACACATCTCTGGCATT); inserts were also checked by PCR with primers that go outwards the insert (F 5’-TCAATGCCAGAGATGTGTATAA R 5’-CACCGCGTAGATGTGT). Samples from the few repetitions were pooled and sent for PacBio sequencing (SciLifeLab, NGI Sweden).

### Analysis of long-read WGS libraries after Tn5 transposition

Alignment was performed using pbmm2 v.1.13.0 (https://github.com/PacificBiosciences/pbmm2), a wrapper for minimap2 (38), against the T2T reference genome, which was indexed using default parameters. Pileups were produced using bamCoverage 3.5.1 from deepTools (39) and visualized using IGV 2.16.0 (40). A custom R script was used to detect the Tn5 insert sequence and perform statistical analyses. Tn5-insert containing reads were gathered and motifs were visualized using ggseqlogo (41).

### Statistical analysis of monocytes

We compared rank(nTA) for the four PBMC batches we had access to (Supplemental Figure 4b-c). If all cells are treated equally, then nTA is different between classical and non-classical monocytes with p < 2.2e16 (t-test). Since multiple batches are present, we also statistically compared the averages. All batches result in p<0.064. Library #4 (pbmc_granulocyte_unsorted_10k) appears to not capture differences in nTA; removing this library would result in p<0.038.

### FACS cell cycle analysis of monocytes

The research was carried out according to The Code of Ethics of the World Medical Association (Declaration of Helsinki), and an ethical permit was obtained from the Swedish Ethical Review authority (#2016/53-31). Blood samples were obtained from human healthy male adults, ages 18-30. Peripheral blood mononuclear cells (PBMCs) were isolated by gradient density centrifugation using Ficoll-Paque PLUS (Cytiva, 17144002) and stored at -150°C. Thawed cells were counted and brought to 10^6^ cells per tube in 100 uL of FACS staining buffer (0.1% sodium azide, 2% FBS in PBS). The cells were then stained for 30 mins at 4°C in the dark with anti-human antibodies specific for anti-CD14 [APC/Cyanine-7] (301819, BioLegend), anti-CD16 [Fluorescein isothiocyanate (FITC)] (360715, BioLegend). For cell cycle analysis, cells were additionally stained by Hoechst 33258 (94403, Sigma-Aldrich) at 10 μg/ml for 30 min at room temperature and additionally stained for viability discrimination by 5 PI μg/ml (P3566, Thermo Fisher Scientific). Samples were washed, resuspended in FACS staining buffer, and taken for FACS analysis. At least 30, 000 gated monocytes were acquired in a 9-color BD FACSMelody flow cytometer (BD Biosciences) using FACSDiva software (BD Biosciences) and analyzed using FlowJo software (TreeStar, v10.9, Ashland, OR). Gating of the monocytes was done excluding the doublets and without the exclusion of dendritic cells. The cell cycle of the monocytes was assessed by fitting a mixture model using the FlowJo Cell Cycle function (42). Gated classical and nonclassical monocytes were sorted in the FACS tubes, pelleted by centrifugation, and imaged at 40× magnification with a Zeiss AxioPlan 2 in the bright field for the morphology.

### General analysis of single-cell data

Unless otherwise stated, all 10x Genomics single-cell ATAC-seq data was aligned using cellranger-atac-2.0.0. Multiome RNA+ATAC-seq datasets were aligned with cellranger-arc 2.0.0. Cell types were annotated according to markers from each respective source article, or annotation files as described for each dataset below. Our custom pipeline Telomemore was then used to count reads for each cell having at least 3 consecutive telomere motifs, as for the previous bulk analysis. Single-cell RNA-seq analysis was done using Seurat (11), and single-cell ATAC-seq analysis using Signac (43). MACS2 was used for peak calling (44). Dimensional reductions were done using UMAP (45). We refer to the provided R source code for the precise details of this analysis.

### Specific analysis of 10x Genomics multiome PBMC/monocyte data

We obtained the 10x PBMC multiome PBMC demo dataset from www.10xgenomics.com (“Fresh Frozen Lymph Node with B-cell Lymphoma (14k sorted nuclei)“, “10k Human PBMCs, Multiome v1.0, Chromium Controller”, “10k Human PBMCs, Multiome v1.0, Chromium X”). Cell types were first predicted using SingleR (46), based on the DICE (47), HPCA (48) and Monaco (49) reference datasets. The final cell type annotation was performed using marker genes, applied to clustering by the Leiden algorithm. The nTA-gene correlations are computed on the clusters referenced in the text.

### Specific analysis of scGET-seq single-cell data

Along with FASTQ files for nTA computation, preprocessed tn5 and tnH count matrices were obtained from ArrayExpress (E-MTAB-10218, E-MTAB-10219, E-MTAB-9651 and E-MTAB-9659) and used for clustering. Routine scATACseq analysis was performed using Seurat/Signac. The ratio of TnH/Tn5 was then calculated and compared with nTA.

### Specific analysis of the Zhang2021 chromatin accessibility atlas

Besides GSE184462, the Zhang2021 atlas further depends on reuse of a pancreas dataset, GSE160472(50), which we included. Privacy protected samples (human heart samples on dbGaP: phs001961, human islet samples on dbGaP: phs002204) were not included. Because the sciATACseq cell barcodes reside in the FASTQ name of the sequencing read, and NCBI SRA strips the read names in their upload, we had to use GEO cloud delivery. Due to the size of the data, we did not perform alignment and *de novo* cluster assignment. Instead, existing cell type annotations were downloaded from https://data.mendeley.com/datasets/yv4fzv6cnm/1 (1B_Cell_metadata.tsv.gz).

To find motifs linked to telomere accessibility, a linear model was set up using Limma (51) with models as shown in the main figure. Motif scores were calculated using ChromVar (9) through ArchR. CISBP was used as the motif database (http://cisbp.ccbr.utoronto.ca/) (52).

### Specific analysis of tonsillar B cell multiome data

Library reads were aligned and aggregated using CellRanger ARC 2.0.0. The classification of TCRs was done using TRUST4 (53). The SNPs of the cells were extracted using cellSNP (54), and assigned to donors using Vireo (55). No batch effects between donors were observed. The Vireo doublet score was used to filter out droplets. Cell types were predicted using SingleR (46), based on the HPCA (48) and Monaco (49) reference datasets. Furthermore, cell labels were transferred using Seurat from a previous single-cell RNA-seq-only tonsillar B cell dataset (56), for qualitative comparison. Major axes (GC/non-GC and naive vs switched) aligned but precise clusters did not align satisfactorily. The dataset clusters, however, had similar topology if annotated based on the same marker genes. We did not find discrete clusters corresponding to “activated” nor “FCRL3^hi^”. Cells not of interest (T cells, dendritic cells) were present but ignored in this study. Especially the marker gene CXCR4^hi^ was used as a marker for the dark zone (DZ) All highly varying genes and motif activities were correlated to nTA using Spearman’s method.

Because it has been reported that GC B cells, compared to naive and memory, are higher in TERT and have on average longer telomeres (57), we also investigated this alternative explanation for AICDA^hi^ cells being nTA^hi^. TERT is however upregulated in rather separate S-phase cells without effect on nTA. Telomeres have also been found to be 1.4kb longer in naive T cells over memory T cells (58); however, assuming an analogy for B cells, we cannot see such a trend on nTA in our data.

## RESULTS

### The telomere proper is protected from transposition, making ATAC-seq unsuitable for telomere length inference

Telomere length has previously been quantified from whole-genome sequencing (WGS) data (24, 26, 59). All current computational methods are based on the classification of reads based on their content of telomeric motifs. The first implementation showed that 3 occurrences of TTAGGG in a read is sufficient to find a correlation with mean length of terminal restriction fragments (mTRF), which is often considered the gold standard for telomere abundance measurement (24). From a WGS library, it is only possible to compute the fraction of telomere-like reads (normalized telomeric abundance). To get actual telomere length, this must be multiplied by a constant fitted from matching mTRF data. A naive estimator of telomere length is thus:

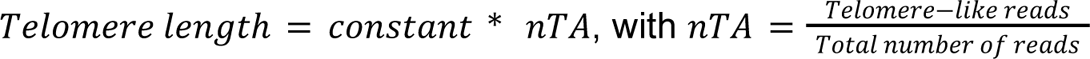

Later methods have slightly improved upon this by normalizing by the amount of telomere variant repeats (TVRs), sequences mainly present in the subtelomeric regions (26). This makes it possible to account for abnormal karyotypes, commonly found in cancer, which have different numbers of telomeric arms. Our initial idea was thus to estimate telomere length similarly, but using ATAC-seq data rather than WGS, as scATAC-seq is easy to perform and such data much more abundant. Because ATAC-seq does not randomly cover the genome, and the absence of knowledge on its ability to target the telomeres, it was however not clear that this is possible.

To evaluate whether ATAC-seq data could be used to infer telomere length or not, we computed nTA for one of the largest bulk ATAC-seq datasets available, from PBMCs of 54 men and 66 women (25) (divided into 3 batches, analyzed separately to account for batch effects). However, unlike previous analyses using WGS data, we did not observe the expected decrease in ATAC telomere motif abundance with age (Figure 1c). Looking for potential technical reasons for the lack of correlation, we found large differences in telomere-like fragment abundances across different ATAC-seq implementations (Figure 1f). However, we were unable to find a correlation between nTA and age (Figure 1d), even when accounting for common ATAC-seq quality control metrics (such as nucleosome signal, and percentage of mitochondrial reads; Supplemental Figure 1a). To further confirm this lack of correlation, we also reanalyzed ATAC-seq of cancer samples from TCGA, comparing them to the telomere length inferred via WGS. We found no correlation between nTA and telomere length, even when accounting for differences in tissue origin (Figure 1e, Supplemental Figure 2).

One possible explanation for the lack of correlation is that the telomere-like reads may not be of telomeric origin. While most of the detected reads had similar sequences and appeared to be tagmented at the same position (Figure 1f), there was still enough sequence variation to perform deduplication (Figure 1g). This does not appear compatible with a telomeric origin, as the telomere is almost perfectly repetitive. However, the large amount of telomere-like repeats in the genome, combined with the use of short read sequencing, prevents detection of the origin by means of alignment.

**Figure 1.**
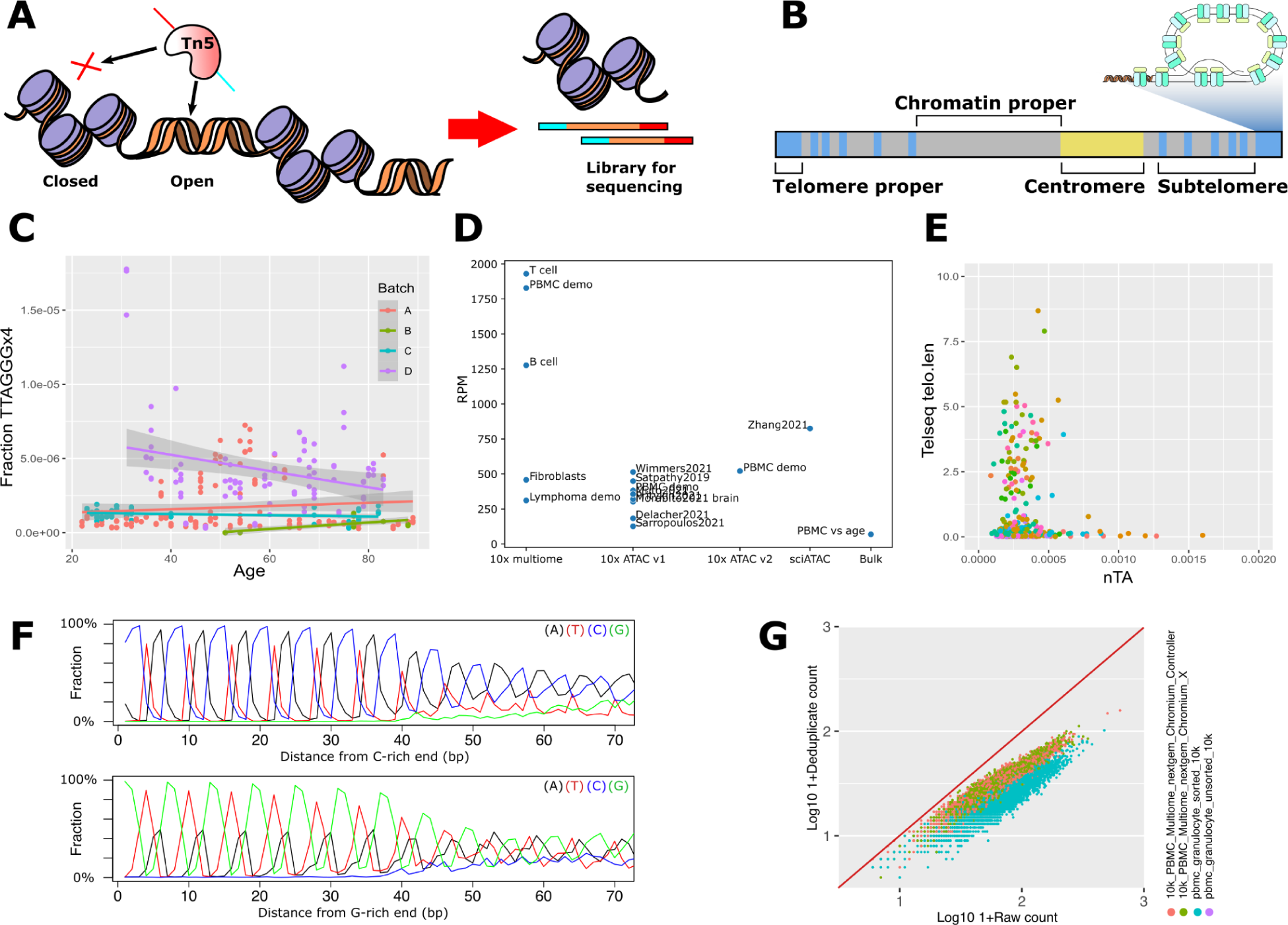
Telomere repeat k-mer-based counting of ATAC-seq libraries does not correlate with telomere length measurement. **(a)** ATAC-seq is a modern method in which accessible chromatin is “tagmented”, i.e. fragmented by the enzyme Tn5, which also adds adapters for sequencing. These fragments are normally used to analyze enhancers. **(b)** Components of the genome, as discussed in the paper. Tandem repeats of TTAGGG are also present in the subtelomere, which does not have a clear end. We refer to the outermost tandem repeat as the “telomere proper”, and what is neither telomere, subtelomere, nor centromere, as “chromatin proper”. **(c)** Normalized abundance of TTAGGG (nTA) across PBMC ATAC-seq datasets does not correlate with age. The common method to infer telomere length from whole-genome sequencing (WGS) data does thus not seem to work for ATAC-seq. The batches A-D are described in methods. **(d)** There is a large variation in nTA across different datasets and ATAC-seq methodologies. The B cell and fibroblast multiome datasets are included in this study; the T cell dataset is in a separate publication (60). **(e)** The ATAC-seq nTA does not show correlation to telomere length in TCGA cancer samples. Coloring by cancer type to show that the type does not appear to influence the correlation (3 outliers cropped; legend in Supplemental Figure 2b). **(f)** Average motif of telomere-like reads, after alignment by GC-content. **(g)** Deduplication shows that motif-containing reads have large sequence diversity.

**Supplemental Figure 1:**
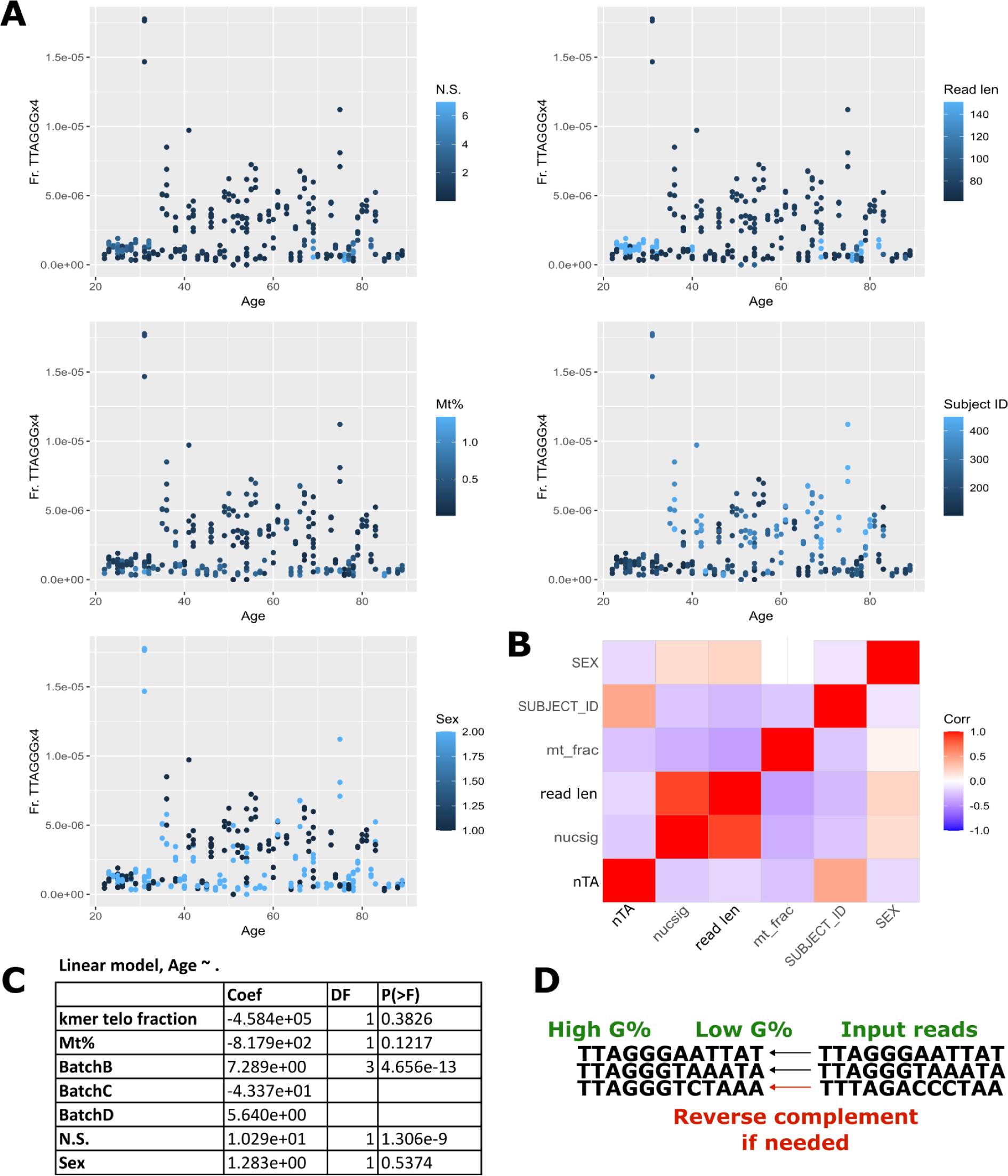
Analysis of confounders in PBMC age prediction. **(a)** Data points colored by possible confounding variables (Nucleosome signal, Mitochondrial % of reads, Sex, Average read length, Subject ID). **(b)** Correlation of QC parameters to telomere-like read counts. **(c)** Example linear model, trying to regress out confounders. **(d)** Reference-free alignment of reads of the purpose of motif construction.

**Supplemental Figure 2:**
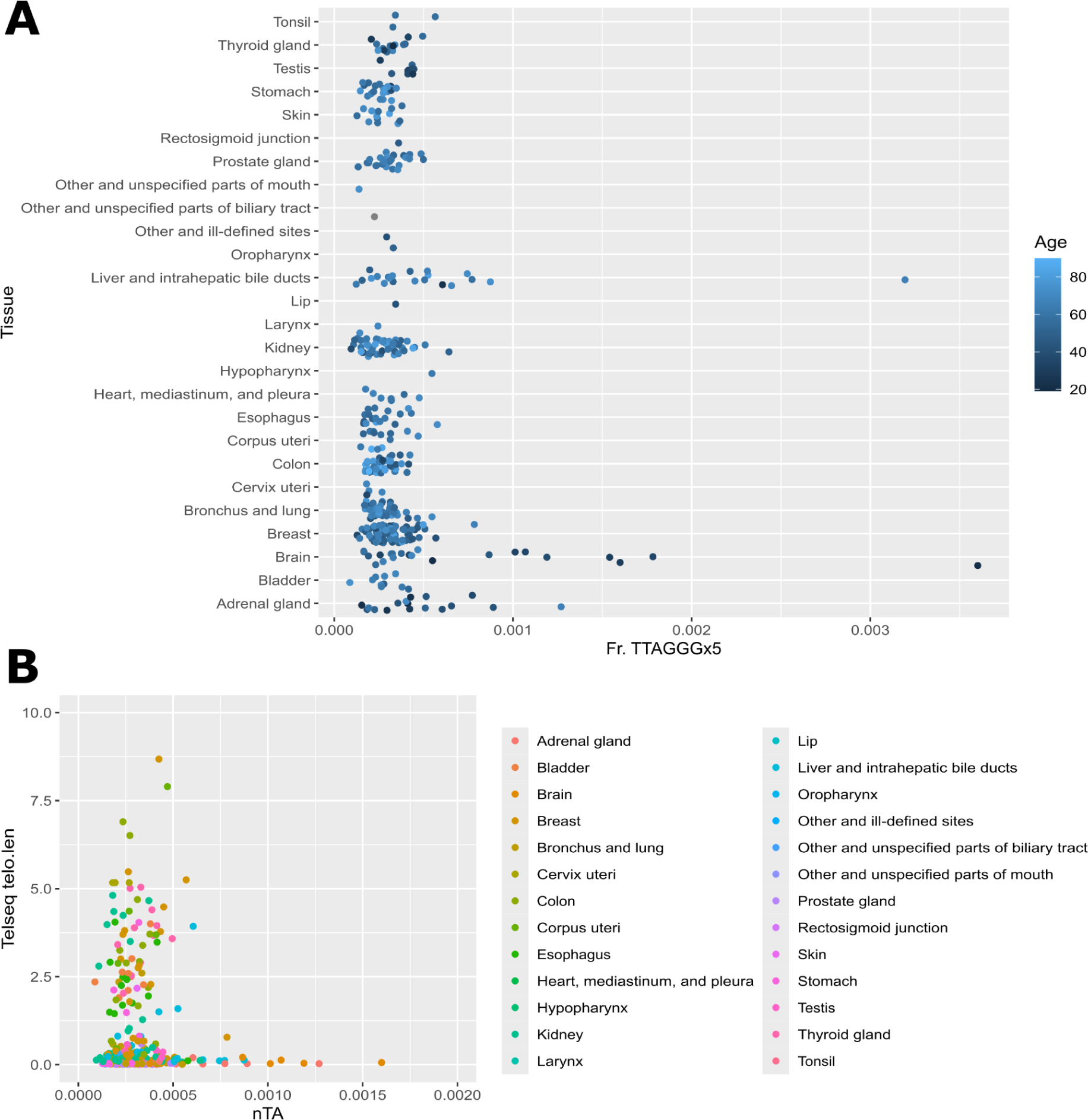
Analysis of confounders in TCGA telomere length prediction. **(a)** Data points separated by tissue origin. **(b)** Scatter plot of nTA vs estimated telomere length, colored by cancer type.

To overcome the limitations of the short-read data, we performed an experiment on two cell lines with known telomere length using long-read sequencing: CCRF (∼7.5kb (61)) and Jurkat (∼4kb (62)). We designed the experiment (Figure 2a) to specifically test if (1) the telomere is hypotagmented, in which case there would be too few fragments to reliably estimate the length; or (2) hypertagmented. In either scenario, fragments would get lost during the ATAC-seq library size selection. Hypertagmentation would further prevent alignment, despite access to long-read sequencing. To overcome these problems, we performed ATAC-seq but with a complete non-fragmenting insert, requiring a separate post-transposition fragmentation. Because Nanopore library preparation itself commonly relies on a transposase, we further opted to use PacBio technology, where the input DNA is mechanically sheared in an unbiased manner. A large amount of cells was used to further avoid the need for PCR.

The result is effectively WGS (∼12X coverage per sample; Figure 2b), but with random Tn5 insertions. Care was taken in the analysis, as we note that Tn5 still incurred some level of fragmentation, likely from some Tn5 homodimers capturing and transposing two separate, rather than one single, DNA insert. Overall, a total of 1.2M Tn5 insertion sites were detected. By computing nTA on all of the reads (i.e. as in TelSeq), Jurkat is predicted to have 14-35% of the telomere length compared to CCRF, somewhat less than their expected telomere lengths (Figure 2c). However, if this analysis is performed only on reads having transpositions, as would be the case for standard, short-read ATAC-seq, this difference disappears (Figure 2d-e). We aligned the reads to the T2T reference genome, thus overcoming the limitations of the GRCh38 reference genome (specifically, that GRCh38 has poor coverage of repeat regions; e.g. an entire 15mb of one arm of chr13 is masked). This alignment verifies the fraction of telomere proper reads with transpositions was lower (Figure 2f). The source of telomere-like motifs is thus primarily from other parts of the genome when performing ATAC-seq.

To understand why the telomere appears poorly tagmented, we performed a motif analysis. It is already known that Tn5 is known to have a preference for GC-rich regions (63). As the telomere motif TTAGGG is rich in G, it is plausible that it is tagmented. The GC bias was confirmed by our per-base motif analysis (Figure 2g). Furthermore, when searching for inserts near or in TTAGGG motifs, we found that Tn5 prefers TTAGGG/ or TTA/GGG (Figure 2h), in line with the motif from short-read data (Figure 1i). This can not be predicted solely by the per-base motif model; however, other factors affect the transposition rate as well, including the higher-order DNA structure (64), and–crucially for ATAC–any bound proteins. The telomere is also known for the presence of G-quadruplexes (65), which may further affect the transposition rate. We thus believe these factors, as part of the shelterin complex, affect the tagmentation process.

While performing the motif analysis, we also note that our per-base motif (Figure 2g-h) does not match well with a previously published motif (66) beyond the GC bias. Most importantly, it has been reported that wild-type Tn5 causes a 9bp duplication of its target (67), but we did not observe this with our hyperactive Tn5_E54K, L372P_ (Figure 2i). It is possible that the mutations responsible for the increased activity (36) have changed this behavior. Thus, while the 9bp duplication is widely assumed in modern ATAC-seq analysis pipelines (43, 68), this may have to be revised, as ATAC-seq is always performed using a modified Tn5.

To conclude, we show that Tn5 has low affinity to the telomere proper during ATAC-seq. This partially explains why any correlation between telomere motif abundance and telomere length likely is too weak to be of practical use. *For brevity, we will use the term “telomere accessibility”, or nTA, to refer to the relative number of reads in ATAC-seq having telomeric motifs, despite that the origin is primarily from the subtelomere*.

**Figure 2.**
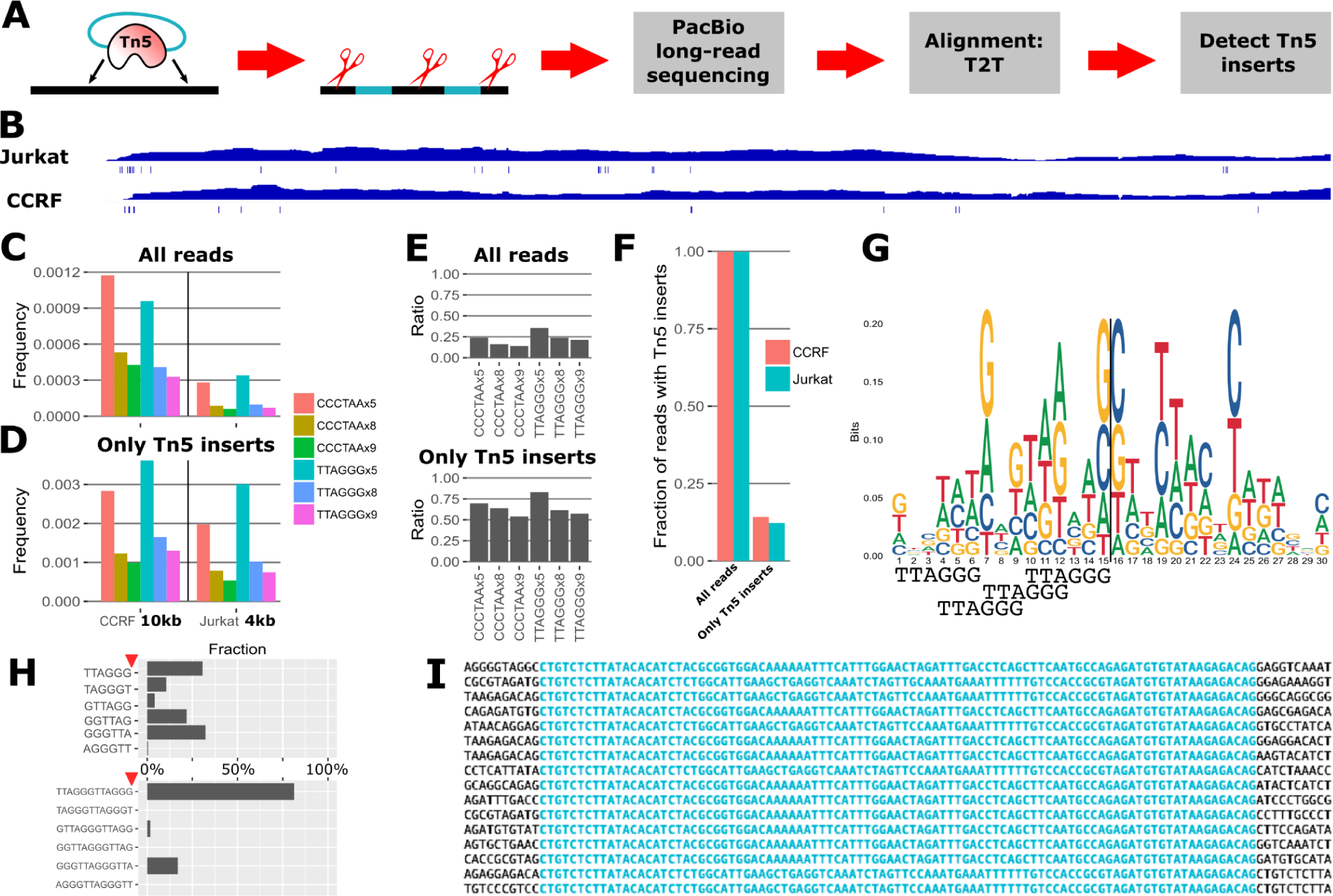
The telomere proper appears to be protected from transposition. **(a)** Experimental setup to find the precise insertion sites of Tn5 during ATAC-seq; Tn5 with non-fragmenting inserts target the genomic DNA, which is then mechanically sheared, and sequenced using PacBio. **(b)** Example coverage of the T2T genome, and detected Tn5 inserts beneath. **(c)** Telomere k-mer-based content can estimate relative telomere lengths for cell lines of known length (i.e. the frequency is much lower in Jurkat than CCRF), **(d)** but not when Tn5-insert-containing reads alone are counted (the frequencies are similar). **(e)** The ratio of nTAs between cell lines (which should be about 0.25) is more correct when searching for shorter telomere repeats, possibly because higher stringency leads to loss of valid reads. **(f)** Fraction of Tn5 insertions in all fragments vs those in the telomere, indicating poor coverage transposition of the telomere proper. **(g)** Per-base motif analysis around transpositions. **(h)** Abundance of telomere motif near transposase inserts, showing a strong location preference that might be further reinforced by chromatin structure. **(i)** No duplication of genomic target sequence is seen, contrary to previous reports (67).

### Differential accessibility of telomere-like repeat regions pinpoint broad chromatin condensation events

To further rule out the possibility of quantifying telomere length, we investigated which other factors influence the nTA measure. We reasoned that the general biological state of the chromatin may impact the readout. The condensation during the cell cycle is an extreme case, which further allows us to compare nTA for the otherwise most homogenous cell population we can imagine. A previous study of DNA accessibility using MNase showed that the S phase is more open than the G_1_ and G_2_ phases, while the M phase is the most condensed (69). We reanalyzed GM12878 cells, FACS-sorted for different cell cycle phases using DAPI, combined with fluorescent Tn5 inserts (“ATAC-see”) (70). It has previously been speculated that early and late G_1_ phase can be separated by the total fluorescent ATAC-see signal, which increases with higher numbers of transpositions (70). This already shows that the cell cycle is an important driver of broad accessibility. The fluorescent signal, however, remains the same across other phases of the cell cycle. Our closer analysis shows that nTA differs little between early *vs* late G_1_, but is markedly low in S-phase (Figure 3a). Aligning the telomere motif-containing reads to the T2T reference, we found that almost all of them map near the subtelomere (Figure 3b), but with rather poor mappability. The signal appears to originate from all chromosomes equally (Figure 3c). This analysis, however, suffers from limitations (i.e. due to the mapping of short reads to repetitive and poorly studied regions). To circumvent this problem, we also assessed similarity between samples using *k*-mer-based Mash distances (71). Telomere-like reads have higher similarity than the genome average (Figure 3d); but the similarity is low (<13%), suggesting a broad origin rather than a single hotspot. The S-phase telomere-like reads stand out as the most distant according to Mash, but possibly because of their small proportion (about a quarter of the amount in other samples). The distribution of the number of telomere-like motifs in reads was not significantly different across samples (Figure 3e).

Because the alignment of reads against short, repetitive regions is not reliable, and because motif-based analysis is rather limited, we also attempted an approach in which we approximately inferred the position of reads using a convolutional neural network. We assumed the genomic model of Figure 1b, but since the subtelomere is poorly defined, we attempted to compute the log(distance) of a read from the telomere (Figure 3f). Furthermore, the chromatin proper and centromere were given artificially high fixed distances (see Methods), reflecting their typical distance from the telomere. As training data, we used the T2T genome sequence for the centromere and chromatin proper. To account for the poorly studied but highly variable subtelomere and telomere regions, we obtain long-read sequencing data across 147 individuals (28). The fitted model was then used to predict the origin of reads for the GM12878 data. As few reads were classified as telomere proper, we binned these with the subtelomeric reads. Overall, we obtained the same results as for the *k*-mer based analysis (Figure 3g), suggesting their equivalence and supporting the idea that the main origin of the telomere-like reads is the subtelomere. As a bonus observation, reads from the centromeric region may be an interesting biomarker for early G_1_ phase. Further study of the centromere is however saved for a future study.

To validate the dependency of nTA on the cell cycle in another cell type, we looked at the orthogonal case of human and mouse naive T cells undergoing activation. Naive T cells, until activated, remain in a condensed dormant G_1_-like state which can even be seen using transmission electron microscopy (TEM) (72). The activation can be seen effectively as synchronized entry into the cell cycle. In line with the GM12878 analysis, we observed that nTA drops during activation as more cells enter S-phase and proliferate (Figure 3h). This estimate is also agreed upon using our machine learning (ML)-based estimate (Supplemental Figure 3).

To conclude, the cell cycle has a large impact on the chromatin condensation state, and by extension, also on nTA. We note similar trends in two cell types. The origin of the telomere-like reads is broad and likely subtelomeric, as supported by alignment, and a novel long-read-informed ML model for approximate alignment. Because accessibility is a relative concept, and ATAC-see FACS showed no difference in total number of transpositions except for early G_1_ (70), we believe a low nTA score is rather due to the rest of the chromatin opening up (Figure 3i). An alternative theory is that the telomeres are replicated at a different time point during the cycle. In yeast, the telomeres are replicated at the end of S phase, which would be compatible; but in mammalian cells, the telomere replication is continuous (73). In either case, it appears possible to use nTA as a biomarker for the cell cycle phase, which is the focus for the remainder of our study.

**Figure 3.**
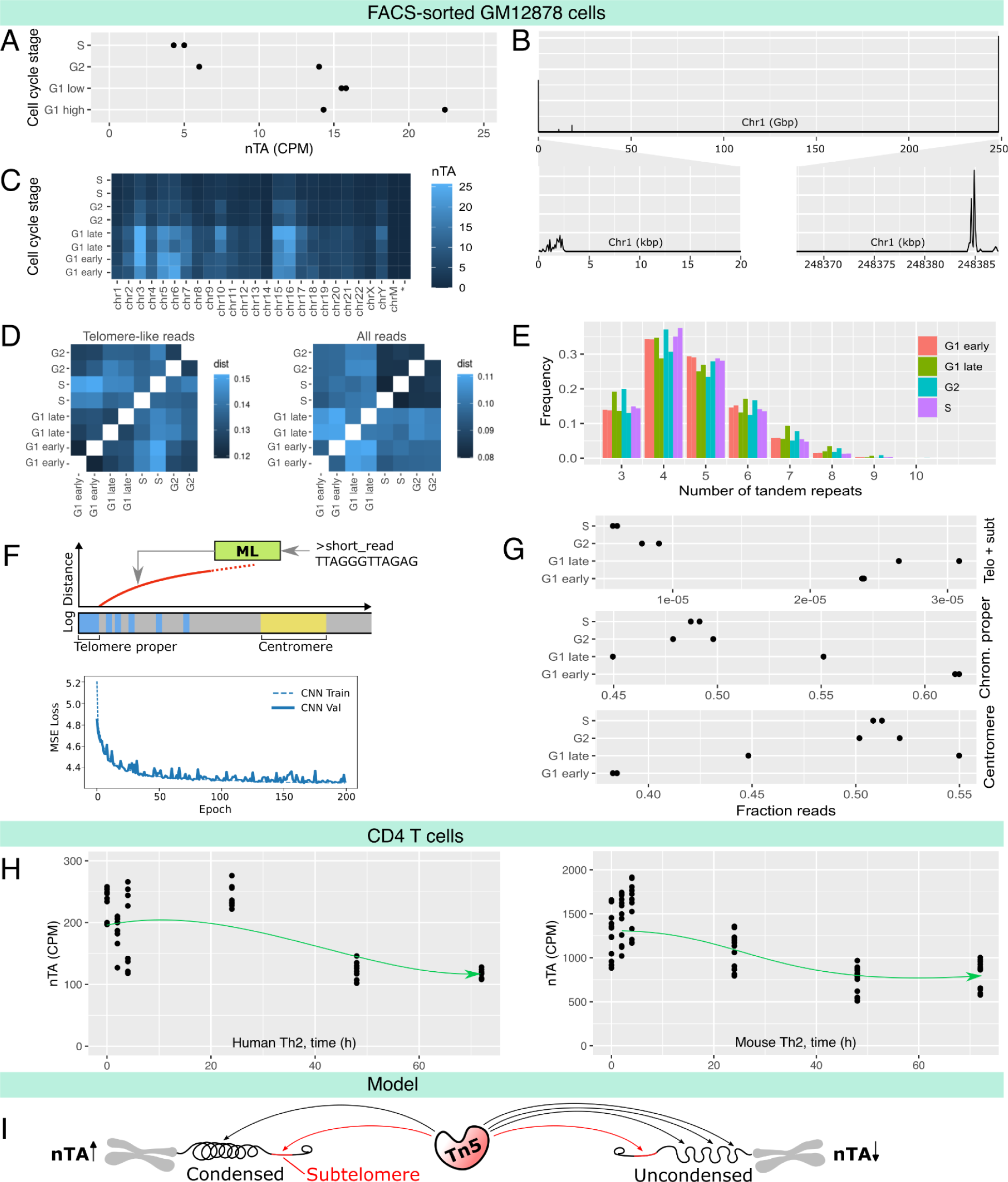
Differential accessibility of telomere-like repeat regions pinpoint broad chromatin condensation events. **(a)** Bulk ATAC-seq nTA from GM12878 cells FACS-sorted by cell cycle stage (70). **(b)** Pileup of telomere-like reads along T2T chromosome 1, suggesting primarily a subtelomeric origin. **(c)** nTA broken down by chromosome, showing that they all co-vary similarly with cell cycle. **(d)** Similarity of bulk ATAC-seq dataset as indicated by Mash and k-mer similarity. **(e)** Cell cycle has little effect on the distribution of telomere motif length. **(f)** A convolutional neural network model to predict the origin of reads. **(g)** Abundance of reads in the different genomic regions vs cell cycle, according to the neural network model. **(h)** Bulk ATAC-seq nTA in human and mouse CD4 T helper type 2 cells during the first 72h of activation, which is a synchronized entry to S phase. The nTA drops as expected (green arrows are conceptual only). Transmission electron microscopy images showing the rapid chromatin decondensation have been generated previously (72)**. (i)** The proposed model of why nTA correlates with chromatin condensation.

**Supplemental Figure 3.**
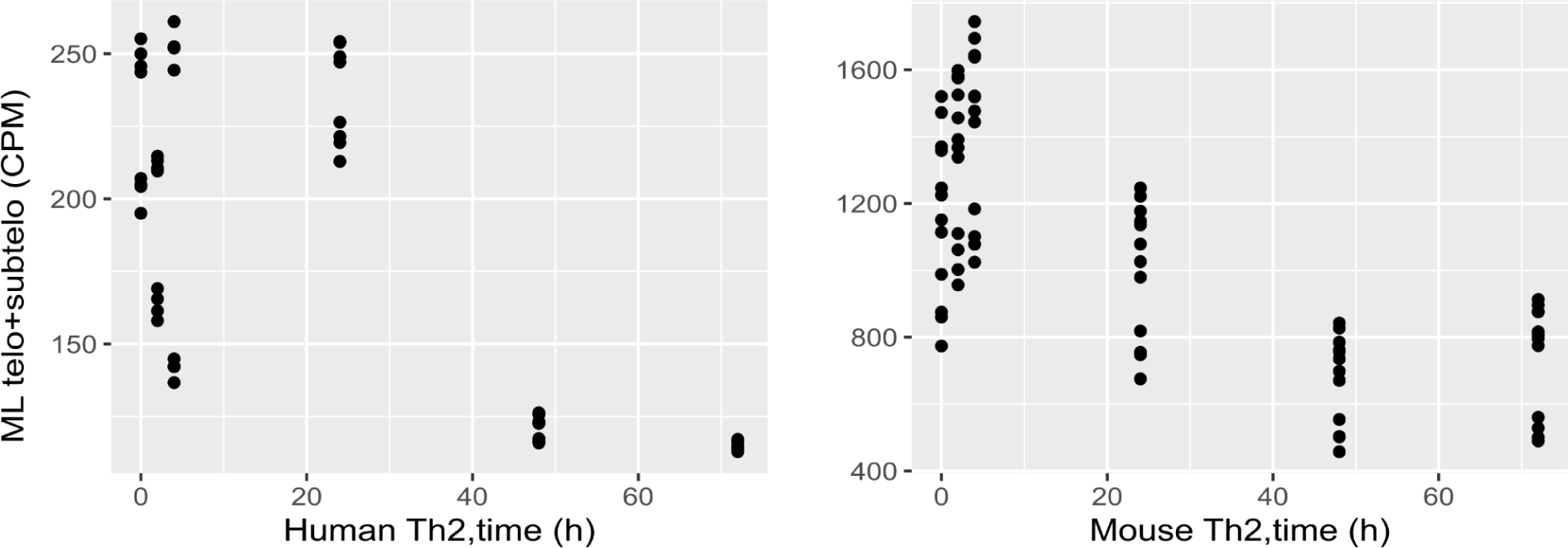
Machine learning (ML)-based analysis of subtelomere abundance in mouse Th2. Bulk ATAC-seq nTA in human and mouse CD4 T helper type 2 cells during the first 72h of activation. The predictions of subtelomere abundance, using the convolutional neural network, are similar to k-mer based estimates.

### Telomere-like read abundance is sufficient for single-cell interpretation of chromatin condensation

To see if nTA can aid in single-cell data interpretation, we focused on two datasets in which the cell cycle is an expected major driver for the clustering. First, we generated a single-cell multiome dataset of primary colonic fibroblasts, as previous RNA-only fibroblast atlases have determined that cell cycle is the major driving component (16). We note that the S case is clearly separated, and that nTA is lowest among these cells, as expected (Figure 4a).

An alternative measure of openness is provided by scGET-seq, a single-cell technology in which normal Tn5 is combined with a modified Tn5 that targets H3K9me3, which decorates the heterochromatin. Thus for each cell, the ratio of open and closed chromatin can be computed, and the ratio is lower in mitotic cells (74). In 3T3 cells, we found a negative correlation between openness (tn5/tnH) and nTA (⍴=-0.12, p<3e-10, Figure 4b), i.e. high openness means low nTA. The previous analysis of this dataset also suggested that the low-openness cells are mitotic (74), in line with nTA being linked to the cell cycle.

To conclude, the cell cycle phase is a strong driver of nTA, and this is especially clear in simple cell types (i.e. where other types of heterogeneity is low). We have, however, assayed most public multiome single-cell data to date (nTA for all datasets available in Supplemental Data), and we have observed other correlations that are harder to untangle. An example is mouse embryogenesis (Supplemental Figure 5), where nTA still correlates with cell cycle, but likely also cell type. nTA must thus be interpreted within the context of other markers to be truly informative.

**Figure 4.**
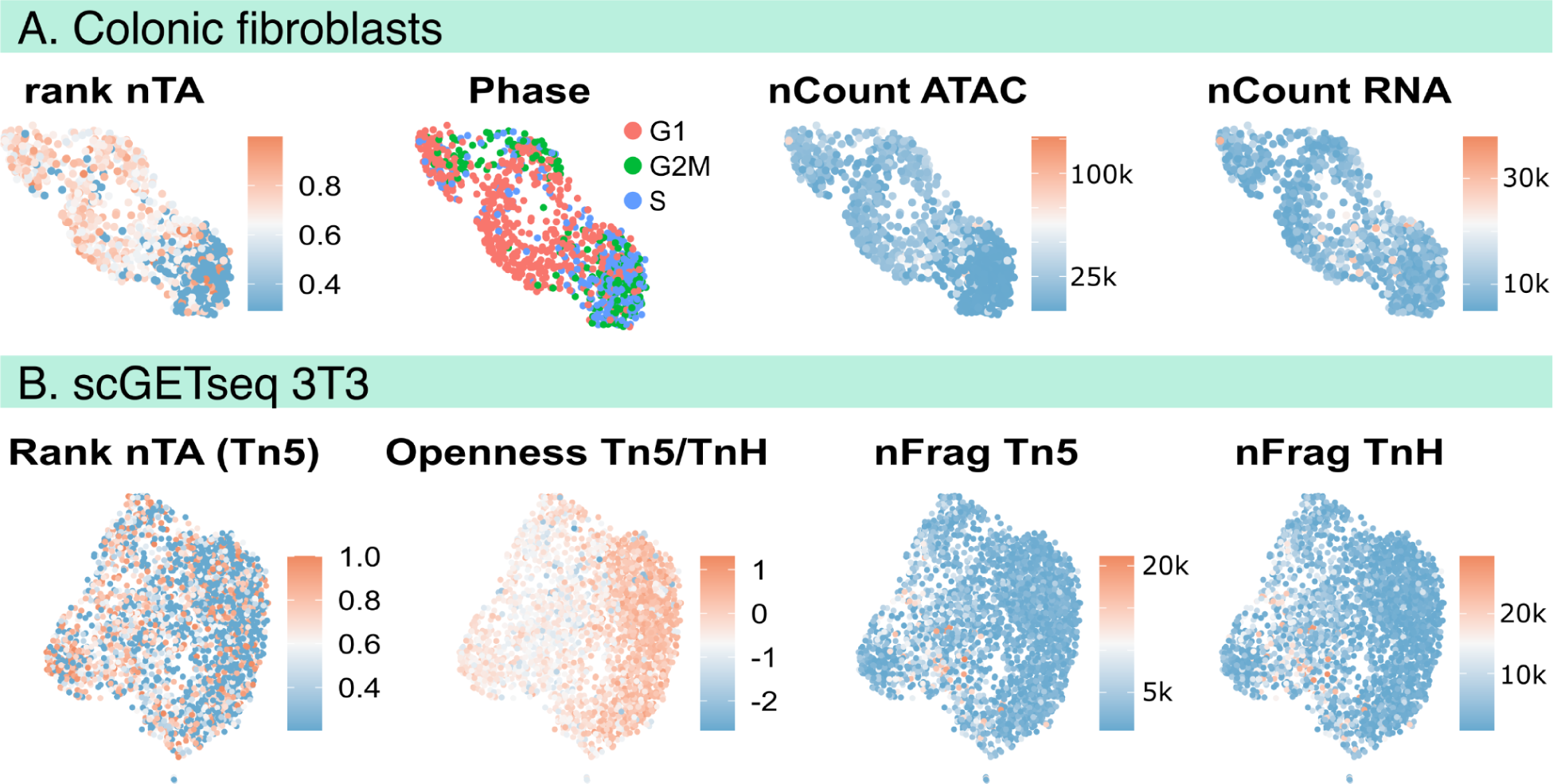
Telomere-like read abundance is sufficient for single-cell interpretation. **(a)** UMAP of a new atlas of primary colonic human fibroblasts cells (one single reaction); nTA is the lowest in S phase, consistent with bulk measurements (Figure 3). Quality control panels indicicate number of ATAC and RNAseq fragments. **(b)** UMAP of 3T3 cells, previously measured by scGET-seq (74); this method measures both open and closed chromatin for direct computation of relative openness, which is negatively correlated with nTA. The panels indicate rank nTA, the openness defined as the ratio of Tn5 to tnH fragments, number of Tn5 and TnH fragments.

**Supplemental Figure 4.**
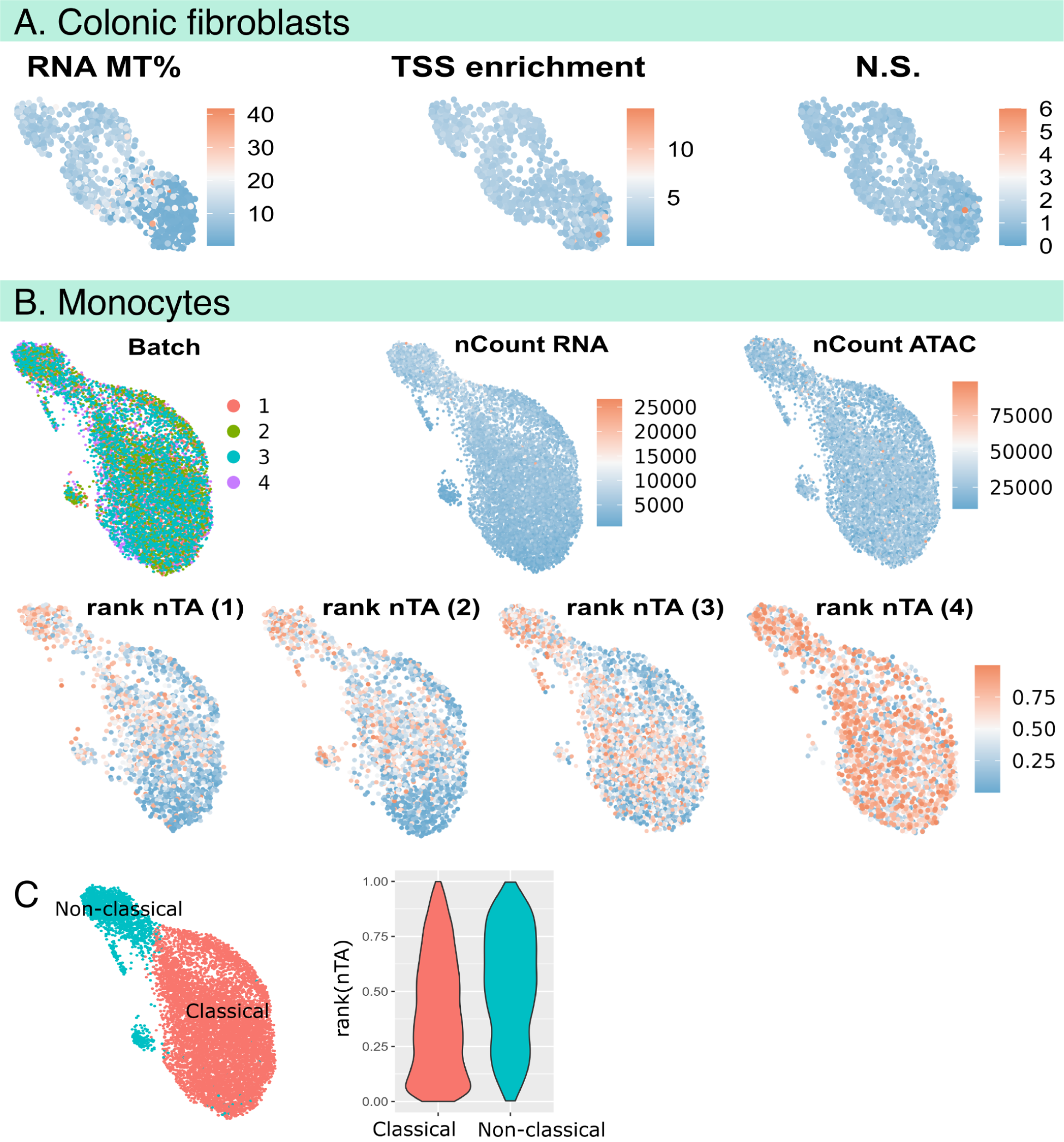
Further quality control measures of multiome datasets. **(a)** Quality metrics for the multiome atlas of primary colonic human fibroblasts. The panels indicate RNA mitochondrial % of reads, TSS enrichment and nucleosome signal. **(b)** Quality metrics for the monocytes in the PBMC multiome atlas, including the mixing of cells across batches, the number of RNAseq and ATACseq reads. Rank nTA is shown for each of the four batches. **(c)** Average nTA for each subtype, with subtypes defined by clustering analysis.

**Supplemental Figure 5.**
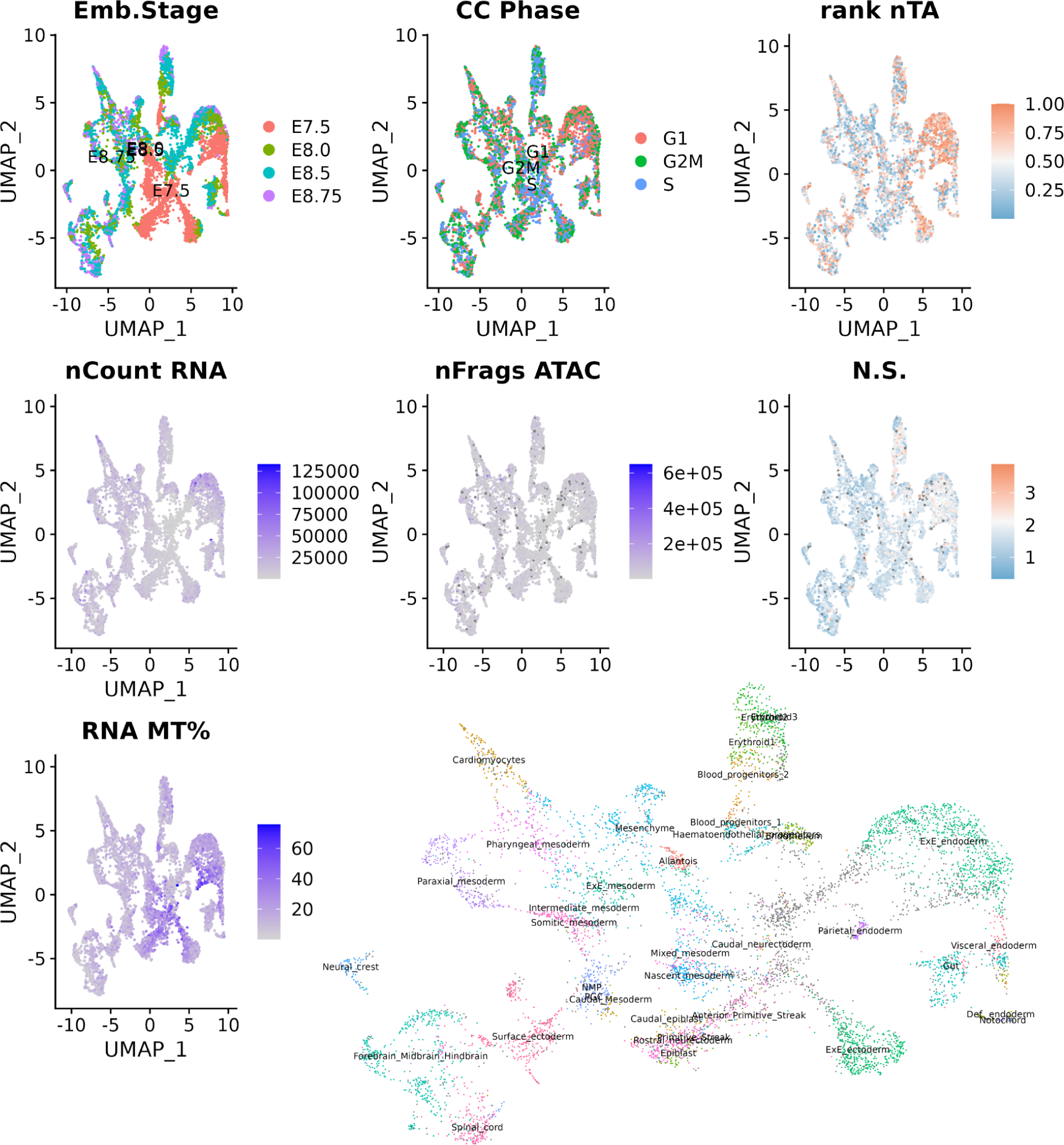
A complex example of nTA in mouse embryogenesis. Multiome data from a previous study (22). There is a strong overlap of G1 phase cells with a region of high nTA, as expected from other results in this study. However, nTA is also high in other locations, suggesting the existence of cell type-specific and other contributions to nTA. Nucleosome signal does not correlate well with nTA and is thus not an equivalent measure. The cells are separated across multiome runs in a non-random manner, making unbiased comparison difficult. Rank nTA is computed per-batch, which will overcorrect differences between the 8 batches, organized by stage.

### nTA complements RNA-based cell cycle analysis

To see if nTA has predictive power, we reanalyzed existing public single-cell multiome data. We found a human PBMC dataset from 10x Genomics (25, 000 cells), which contains monocytes. These are especially interesting from a chromatin condensation perspective, as they are generally characterized by bilobed, horseshoe-shaped nuclei, which we figured might have a strong impact on chromatin accessibility patterns. However, there are multiple monocyte subsets, with different functions (75). In this single-cell dataset, non-classical CD14^lo^CD16+ monocytes have a higher nTA (Figure 5a, quality metrics in Supplemental Figure 4b). A potential explanation is that the second-most positively nTA-correlated gene is CDKN1C (24%), which inhibits proliferation during the G_1_ phase (76). We verified using FACS that this subset indeed is stalled in G_1_ (Figure 5b-c). Interestingly, this is in disagreement with typical RNA-seq-based cell cycle annotation (Figure 5a). Furthermore, the subsets are clearly different morphologically, with non-classical monocytes having less granules (Figure 5c).

This example highlights the difficulty in precisely annotating the cell cycle phase from RNA-seq data–notably, at least 16 different bioinformatics tools/methods exist for this purpose to date (77). While we do not claim better performance, we do however postulate that ATAC-seq provides orthogonal data, and that chromatin condensation/decondensation needs to happen independently of which genes are transcriptionally regulating the process. Thus, multiome analysis might be a better choice for applications when the cell cycle state is of particular interest.

**Figure 5.**
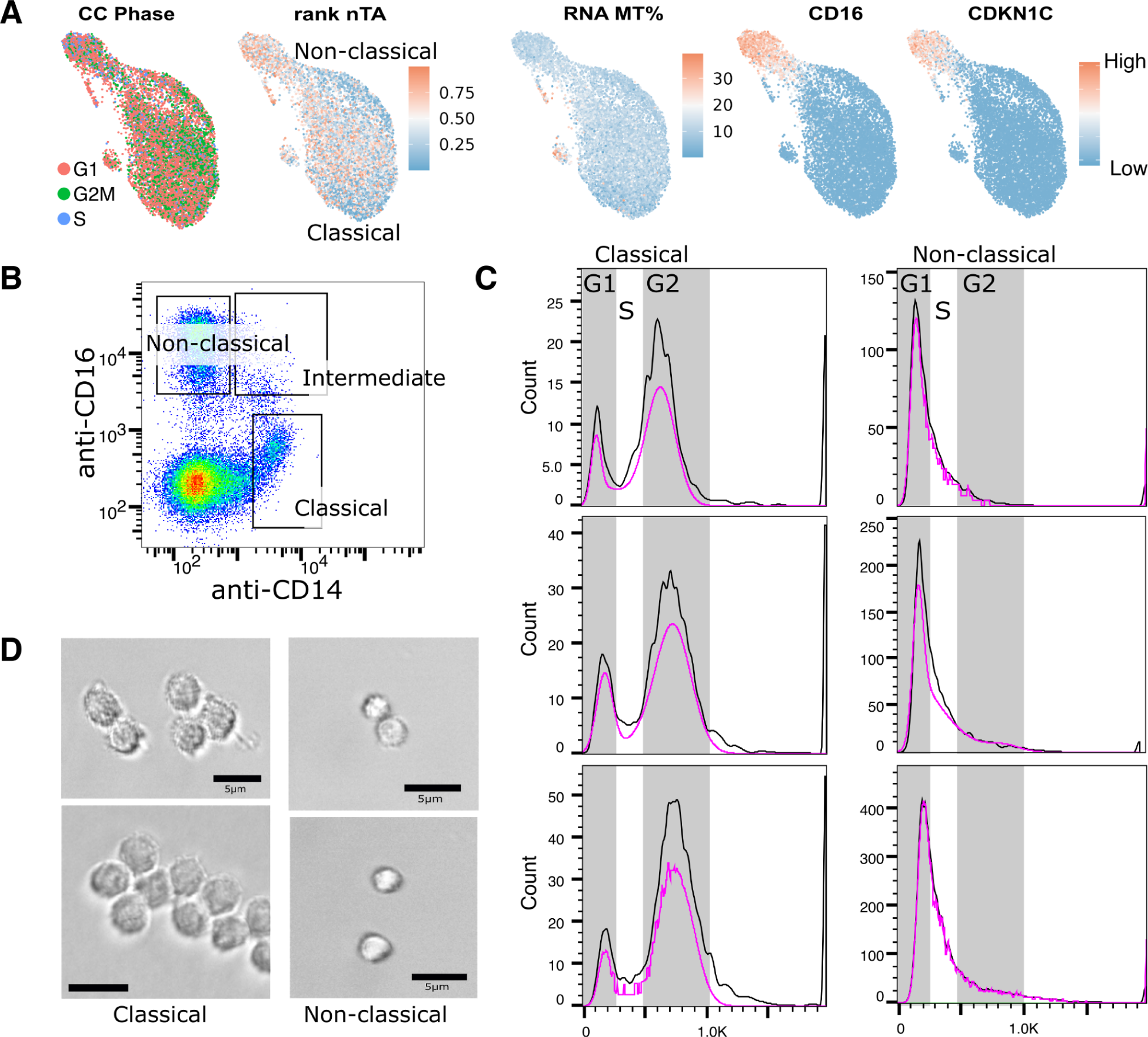
Comparison of monocyte telomere accessibility vs cell cycle. **(a)** UMAP of monocyte cells in the 10x PBMC multiome dataset, showing the nTA, markers for subsets, and RNA-based prediction of cell cycle phase. Based on the possible elevation of nTA in the non-classical subset, we performed further analysis of condensation. CDKN1C is one of the most correlating genes with nTA that is likely to explain condensation in terms of cell cycle. **(b)** FACS gating for the sorting of different monocyte populations. **(c)** DAPI staining of monocyte populations to determine cell cycle phase distribution. **(d)** Images of monocyte populations, where classical monocytes appear bigger and more grainy than the non-classical subset.

### ATAC-seq across tissues and cell types pinpoints cell cycle-related transcription factors

As we expected nTA to also be driven by cell type-specific factors, and not just the cell cycle, we reanalyzed a recent sci-ATAC-seq atlas of chromatin accessibility in the human genome (21). The nTA of the publicly available subset (500k cells) is shown for different cells across tissues in (Figure 6a, nTA values in Supplemental File 1). Naive T cells have a high nTA, reflecting previous observations of their condensed state. To find the most common drivers of nTA, we set up two linear models to infer nTA from TF motif activities. By including a large number of cell types at once, we expected that most cell type-specific effects would be averaged out. The results were similar, with and without including cell type-specific baselines in the model (Figure 6b-c, correlations in Supplemental File 2). At the global level, the activity of motifs TP63, GRHL1 and FOSL2 are among the most positively correlating with nTA. Notably, these motifs have links to the cell cycle. Among the TP transcription factor family members, which have similar motifs, the tumor suppressor and cell cycle inhibitor p53 is arguably the most famous. Knock-out of GRHL1 also leads to cell cycle arrest in G_2_/M phase (78), this most condensed phase. Conversely, AR, MEF2D and NR3C1 are negatively correlated with nTA. The androgen receptor (AR) is important for breast and prostate cancer proliferation (79). The pleiotropic MEFs are activated at the G_0_/G_1_ transition, and promote S-phase entry (80).

Overall, global TF activity inference detects TF motifs correlating with nTA, which are related to cell cycle regulation and are consistent with the interpretation of nTA as a marker for chromatin condensation. We believe further interesting correlations can be found in individual datasets; the computed nTAs are thus available as a resource in the Supplemental Data for further exploration of cell type-specific drivers.

**Figure 6.**
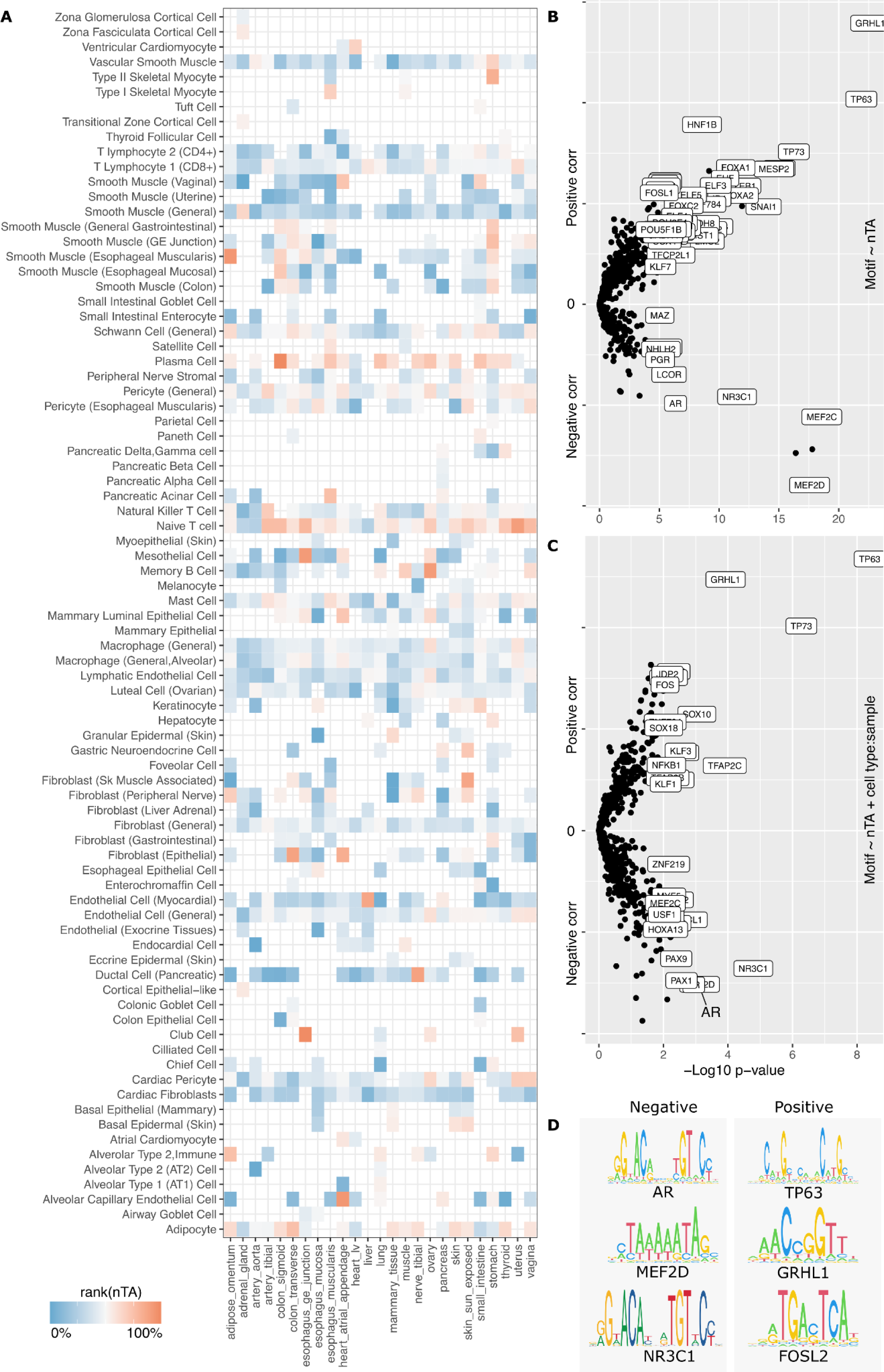
A single-cell atlas of telomere accessibility in the human genome. **(a)** Normalized telomere accessibility across human cells and tissues (21). **(b)** Linear model showing motifs of enhancers that are correlated with nTA. **(c)** Linear model also removing cell type-specific differences. **(d)** Motif sequences of some of the top drivers of nTA.

### ATAC-seq implicates putative drivers of chromatin condensation during B cell somatic hypermutation

As an example application of how the nTA measure can be used for hypothesis generation in a typical single-cell setting, we generated and applied our measure to an atlas of human tonsillar B cells across 5 individuals (Figure 7a-b). B cells are the source of antibodies, which evolve in the lymph node germinal center through two modes of genome self-editing: somatic hypermutation (SHM) and class switch recombination (CSR). As off-target mutations can lead to cancer, this process is tightly controlled (81). It has been shown through microscopy that the mutagenic enzyme AICDA is present in cytoplasm and only enters the nucleus during cell division, likely as a titration mechanism. Thus, mutagenesis happens during early G_1_ phase (82). We speculate that the simultaneous chromatin condensation also promotes selective mutagenesis of the B cell receptor, and that single-cell analysis of the condensation process may aid in finding regulators of this process.

Our atlas covers both follicular and germinal center (GC) B cells (Figure 7a-b). For the purpose of this analysis, we however focused on the GC B cells only, where SHM/CSR occurs (Figure 7c-d). These cells are primarily separated along the light/dark zone gradient (CD83-CXCR4), but also by cell cycle. This is in line with classical models of SHM/CSR and expansion happening in the dark zone (83). The gene AICDA, responsible for SHM/CSR, is especially expressed among cells in the G2/M phase, extending into G1 phase. Similarly, nTA does not align with the RNA-based cell cycle annotation, but has a similar pattern (Figure 7c, e). Overall, while we cannot exclude other drivers of nTA than condensation, these observations are all consistent with the observation that AICDA acts in early G_1_, when the chromatin is condensed (82), and thus must be expressed at some point before G_1_ (i.e. G_2_/M). The level of nTA is consistent with this idea and may thus plausibly reflect the level of condensation.

Based on this assumption, we used correlation to find putative drivers of condensation (Supplemental File 3). The top-most correlating genes are the TF FOXP1 (25%, p<2.5e-24), followed by the TF ZNF831 (25%, p<1.9e-23) (Supplemental Figure 7). FOXP1 has previously been linked to B cell lymphoma (84) while ZNF831 is overall poorly studied. Other correlating games include the actin polymerization regulator KANK1 (22%, p<1.1e-20, Figure 7e), for which KO can lead to cell proliferation (85). nTA also correlates with CDK13 (23%, p<8.6e-20, Figure 7e), which has been shown to increase Pol II processivity (86). GWAS has associated CDK13 with the amount of IgD+CD38^dim^, and IgD+CD24-B cells (87), or lymphocyte count in general (88, 89). It is therefore possible that CDK13 upregulates DNA damage response genes, as previously shown for CDK12 (90), and thereby helps control the dose of SHM. Further validation of these genes is however outside the scope of this manuscript.

Overall, our analysis shows that nTA, for a specific cell type, might reflect known chromatin dynamics. As in the monocyte example, the RNA-based cell cycle annotation might not be the best option, especially if the chromatin state is of primary interest. nTA can then be used to infer putative regulators, such as CDK13, which can aid in understanding how lymphoma may form due to AICDA off-target mutations.

**Figure 7.**
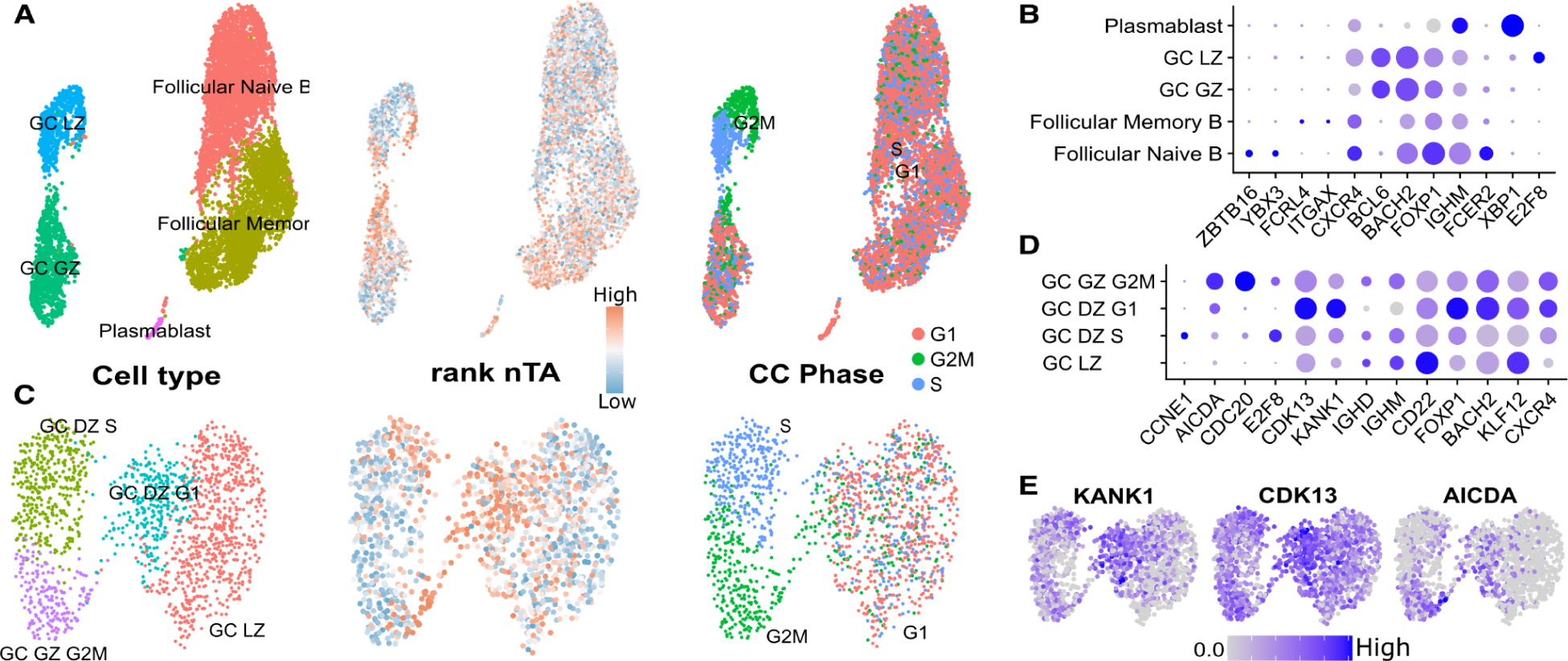
A multiome atlas of tonsillar human B cells. **(a)** UMAP of all B cells. **(b)** Expression levels of some marker genes for all B cells. **(c)** UMAP of GC B cells only. **(d)** Expression level of some marker genes for the GC B cells. **(e)** Expression of AICDA and some of the genes most correlating with nTA.

**Supplemental Figure 6:**
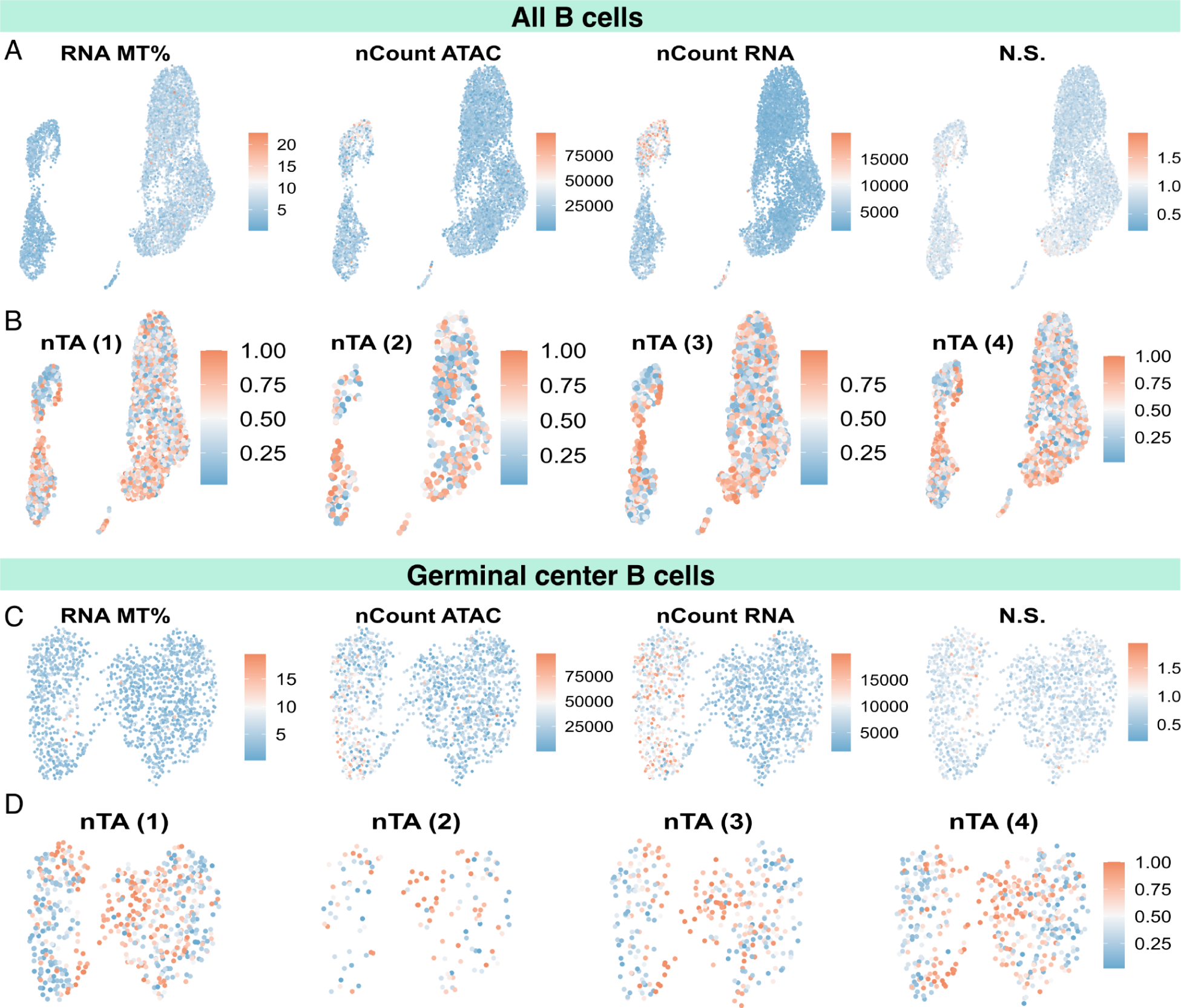
Quality control of the B cell atlas. **(a)** General QC measures for all B cells. **(b)** nTA per donor **(c)** General QC measures for the germinal center cells in particular. **(d)** nTA per donor for the germinal center B cells.

**Supplemental Figure 7.**
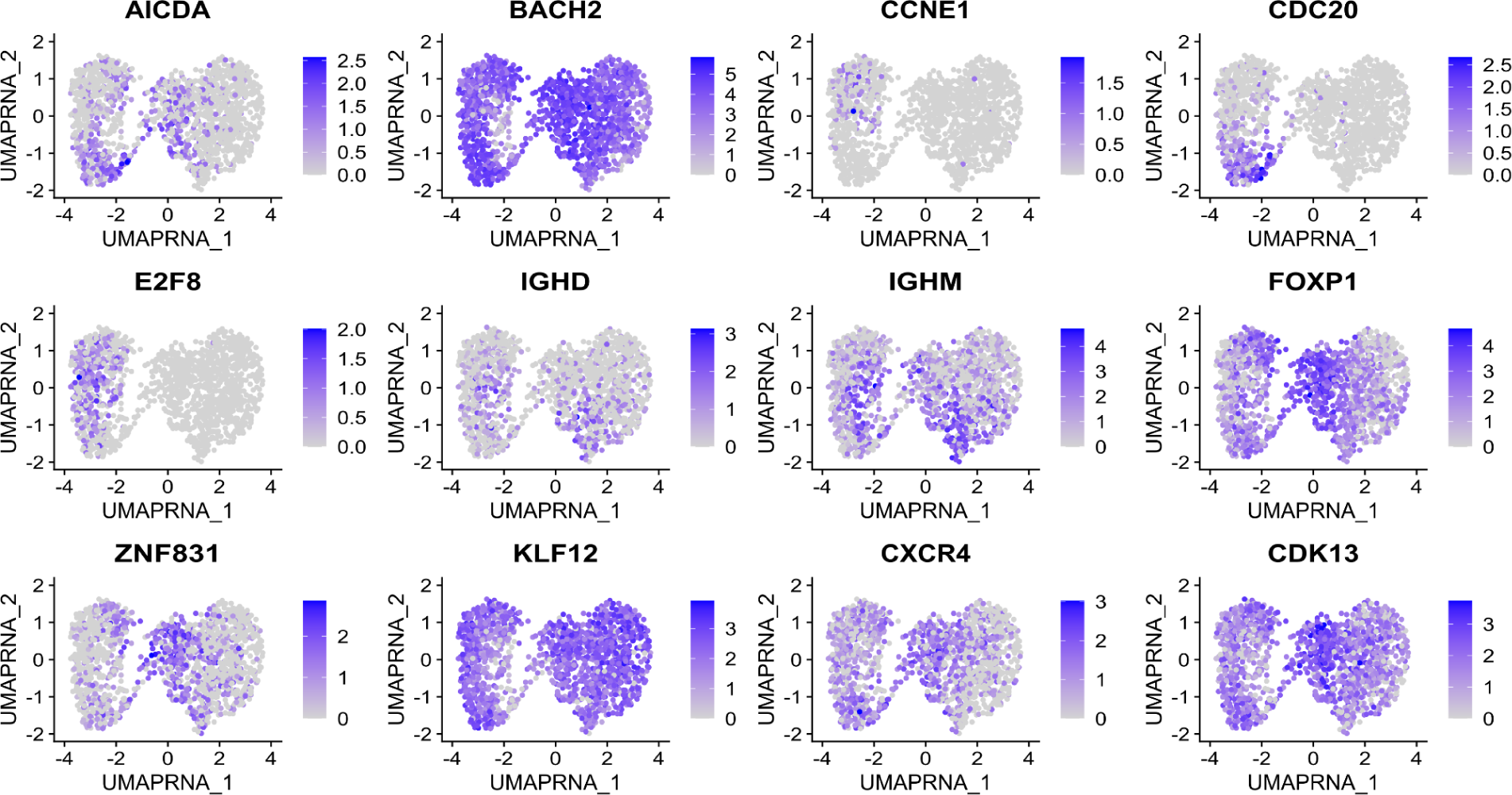
Detailed expression patterns of some germinal center B cell marker genes.

## DISCUSSION

In this study, we aimed to develop a way to infer telomere length from the presence to telomere-like reads in ATAC-seq data, similar to previous WGS-based approaches. By generating a long-read atlas of transpositions, we conclude that the telomere appears largely protected from transposition, resulting in a signal too weak for detection. As it might still be possible to improve the signal-to-noise by accounting for technical confounding factors, we set out to find such confounders. We find the cell cycle to be especially relevant, as differential accessibility to the subtelomere *vs* the remainder of the genome affects the normalized telomere abundance score (nTA). Other quality metrics for ATAC-seq are also affected, such as reads/cell, or nucleosome signal, but we do not find consistent correlations that can be used for compensation in, e.g., a linear model for telomere length. This could otherwise account for batch differences caused by for example different tagmentation times and SPRI bead cleaning ratios (size section). Based on our exhaustive analysis, telomere length prediction at the single-cell or bulk level is not viable from ATAC-seq.

The interpretation of nTA is still somewhat complicated in that the subtelomere is poorly defined; it is rather a continuum that comprises about 25% of the most distal 500kb and 80% of the most distal 100kb in human DNA (91). Telomeres are also of different conserved lengths for each chromosome (28), suggesting that each arm is regulated differently. Here, we only consider the average subtelomeric read, being limited by short-read sequencing technology. Thus, nTA is a crude biomarker, which likely can be refined in future work. Our alternative approach (i.e., predicting the origin of reads using convolutional neural networks) is promising, but requires higher quality training data to reach its full potential.

Accepting that the nTA score is primarily linked to chromatin condensation, in a cell type-specific manner, we however show that it can still be used to help interpret single-cell data. We provide a tool, Telomemore, for the single-cell quantification of mainly-subtelomeric *k*-mer-containing reads. Using monocytes as an example, we find that RNA-seq to infer cell cycle phase is not always reliable. However, it appears that nTA can be used as a complement for data interpretation. Using a large number of different cell types, we further find TFs relevant for chromatin condensation–notably (but not limited to) cell cycle regulators. Finally, we use the nTA measure to study the closing down of the chromatin during B cell somatic hypermutation, which is crucial to avoid off-target mutations and B cell lymphoma. Correlation with nTA suggests that the cell cycle regulator CDK13 may be an important driver for this process.

Further interesting observations were made during this study; notably, the wide assumption that Tn5 duplicates 9bp of the target region appears wrong, at least for the mutated hyperactive Tn5 used for ATAC-seq. ATAC-seq reads of the centromere may also be informative to pinpoint the early G_1_ phase, but more work is needed to validate this correlation.

The production of this manuscript highlighted the need for the FAIR principles (Findability, Accessibility, Interoperability, and Reuse). Much single-cell data has only been made available as processed count tables, with the implicit assumption that it contains all interesting information. However, others have made use of mtDNA to perform lineage tracing (92), and we provide further uses of the raw data. We thus urge others to always release their sequencing data as raw FASTQ files, and ideally also deposit it in the Human Cell Atlas to aid reprocessing. Finally, this study would not have been possible to conduct without easy access to primary data, and a significant amount of effort went into obtaining data access agreements. Thus, we call for a discussion of which data should be considered “sensitive”, as needlessly hiding raw data slows down research and is against the interest of those benefiting from drugs derived from the analysis of human single-cell data.

## DATA AVAILABILITY

All sequencing data generated in this study has been uploaded to ArrayExpress (human tonsillar multiome B cell atlas at E-MTAB-12632, multiome fibroblast atlas at E-MTAB-14220, long-read transposed DNA at E-MTAB-14238). Telomemore is open-source and freely available at Github, https://github.com/henriksson-lab/telomemore.java. Scripts to reanalyze the various datasets, and precomputed abundances, are available at https://github.com/henriksson-lab/telomemore_supplemental. The human tonsil B cell data can be viewed interactively at http://data.henlab.org/.

## SUPPLEMENTARY DATA

Supplemental File #1. Zhang2021 atlas nTA per cell type and dataset

Supplemental File #2. Zhang2021 atlas nTA correlations with TF activities

Supplemental File #3. Correlations of genes to nTA in germinal center B cells

## AUTHOR CONTRIBUTIONS

W.R. conceived the name and implemented the first version of Telomemore. A.D. and R.G. contributed material and helped with flow cytometry. I.S.M. and M.S. performed the single-cell data generation. I.Y. performed the long-read transposon insertion analysis, and validation of the monocyte cell cycle state. J.T., M.F. and J.H. supervised. J.H. conceived the study and developed the Telomemore algorithm and neural network-based inference of approximate sequencing read origin, with input from L.M.C., J.H., I.S.M., M.S., I.Y., and L.M.C. co-wrote the manuscript with input from all co-authors. All authors have read and agreed to the published version of the manuscript.

## ACKNOWLEDGEMENTS

All authors have read and agreed to the published version of the manuscript. We thank Matthew Weirauch for excellent help with the CIS-BP database; Jacob Scheurich at the EMBL Protein Expression and Purification Core Facility for the production of recombinant Tn5_E54K, L372P_; Magnus Hultdin, Sofie Degerman, Pär Larsson and Martin Gullberg (Umeå university) for discussions about telomeres and cell cycle; Xi Chen (SUSTech) about ATAC-seq; and Javier Avila-Carino and Teresa Frisan for help with cell culture. The analysis was inspired by a collaboration with Oliver Billker and Ágnes Regős on malaria ATAC-seq. Filippe Seiz De Filippi did a first attempt at annotating the B cells.

